# Bioprinting the Tumor Microenvironment with an Upgraded Consumer Stereolithographic 3D Printer

**DOI:** 10.1101/2021.12.30.474546

**Authors:** Louise Breideband, Kaja N. Wächtershäuser, Levin Hafa, Konstantin Wieland, Achilleas Frangakis, Ernst H. K. Stelzer, Francesco Pampaloni

## Abstract

A widespread application of three-dimensional (3D) bioprinting in basic and translational research requires the accessibility to affordable printers able to produce physiologically relevant tissue models. To facilitate the use of bioprinting as a standard technique in biology, an open-source device based on a consumer-grade 3D stereolithographic (SL) printer was developed. This SL bioprinter can produce complex constructs that preserve cell viability and recapitulate the physiology of tissues. The detailed documentation of the modifications apported to the printer as well as a throughout performance analysis allow for a straightforward adoption of the device in other labs and its customization for specific applications. Given the low cost, several modified bioprinters could be simultaneously operated for a highly parallelized tissue production.

To showcase the capability of the bioprinter, we produced constructs consisting of patient-derived cholangiocarcinoma organoids encapsulated in a gelatin methacrylate (GelMA)/polyethylene glycol diacrylate (PEGDA) hydrogel. A thorough characterization of different GelMA/PEGDA ratios revealed that the mechanical properties of the bioprinted tumor model can be accurately fine-tuned to mimic a specific tumor micro-environment. Immunofluorescence and gene expression analyses of tumor markers confirmed that the bioprinted synthetic hydrogel provides a flexible and adequate replacement of animal-derived reconstituted extracellular matrix.

## 1. Introduction

In recent years tissue engineering has advanced as a method to improve regenerative medicine and disease modelling. Combining the use of biomaterials as scaffolds and cells, different methods can be used to assemble functional tissues.^[1]^ One of those methods is three-dimensional (3D) bioprinting, which is the accurate deposition of material according to a computer-generated design, or computer-aided design (CAD).^[2]^ 3D bioprinting has been put on the forefront of tissue engineering research as a method to automate scaffold deposition and thus reduce error while increasing resolution. The level of customization available to the users^[3, 4]^ (depending on the resolution achieved by both the machine and the biomaterial used) allows for a broad range of applications, from drug screening^[5]^ to organ transplantation.^[6]^

The potential of 3D bioprinting becomes apparent as a mean to bridge the gap between cell culture and human tissue. This gap is currently largely bridged by using animal models. Indeed, animal models are time-consuming, expensive, ethically controversial^[7]^ and are in many cases inadequate to accurately represent the human physiology.^[8, 9]^ However, the alternative to animal model, two dimensional (2D) and 3D cell culture, are still far from mimicking organ physiology. They either lack three-dimensional organization of cells (in the case of 2D cell culture)^[10–15]^ or are limited, for example by the absence of vascularization or by the variability between different batches of scaffold (in the case of 3D cell culture).^[16]^ Conversely, bioprinting involves highly customizable designs and biomaterials that allow for the modelling of virtually any organ or tissue in the human body in a reproducible fashion.^[17]^ By using functionalized synthetic bioinks, constructs are made safer for implantation in the human body^[18]^ while vascularization lowers chances for graft rejection^[19]^ and simulates the microenvironment of tissues more accurately. For these reasons, tissue engineering with 3D bioprinting at its forefront is a relevant method that could account for some of the drawbacks in other systems. However, the availability of bioprinters to small and medium-sized laboratories is limited by the high cost associated with commercially available devices. Low-cost systems require an investment of minimum $1,500 and high-end systems reach up to $1 million^[20]^. Additionally, the users of commercial bioprinters often depend on the manufacturer for any add-on, customization, maintenance and bioinks. To make bioprinters more accessible, researchers have developed do-it-yourself (DIY) 3D bioprinters.^[21–30]^ Yet, DIY bioprinter are mainly based on extrusion, the process of depositing material using a nozzle continuously extruding material.^[31]^ Those printers either require viscous bioinks exposing cells to shear stress^[32]^, or require a lot of expertise by the users to assemble the system themselves.

With the aim of circumventing some of these limitations, we developed a device based on light stereolithography (SL), customized from a commercially available low-cost and small footprint 3D SL printer. Stereolithography is defined as a process in which “photocrosslinkable hydrogels are selectively solidified in a layer-by-layer manner that additively builds up 3D structure”.^[33]^ SL allows for fast bioprinting as the deposition and crosslinking using light occur simultaneously. It also tolerates a wider range of viscosity among bioinks and does not expose the cells to shear stress, resulting in higher cell viability.^[34]^

We tested the capacity of the customized bioprinter to produce three-dimensional constructs using hydrogel. For this purpose, a mixture of gelatin methacrylate (GelMA) with polyethylene glycol diacrylate (PEGDA) at different concentrations was used to model the extracellular microenvironment of patient-derived cholangiocarcinoma organoids (CCAO). The results of this work show that 1) an affordable consumer SL printer can be straightforwardly modified for bioprinting, 2) the constructs obtained with the modified SL printer shows a high viability of patient-derived cells, 3) the bioprinted cholangiocarcinoma constructs are equivalent to the established Matrigel culture and are thus suitable for basic research and screening assays. We hope that our DIY SL bioprinter will pave the way for a widespread adoption of bioprinting in cell biology and translational research laboratories.

## 2. A commercial three-dimensional printer can be adapted into a versatile 3D bioprinter

A commercial SL 3D printer (Anycubic Photon S) was selected as a readily available and affordable device. It includes a 115 mm x 65 mm printing platform (on which the constructs attach) and a 178 mm x 120 mm vat with a transparent bottom that contains the photosensitive resin. A 405 nm LED array back-illuminates a liquid crystal display (LCD) that produces a slice-by-slice pattern determined by the CAD file, sequentially polymerizing layers of the photosensitive resin. The Photon S device can print objects as large as 115 mm x 65 mm x 155 mm. However, for the purpose of this work, eight smaller objects were printed (up to 4.5 mm x 4.5 mm x 6 mm) to conserve material (cells and medium) while having a high number of replicates. The SL printer was modified to be used as a 3D bioprinter as follows.

First, temperature and CO_2_ controls were installed by drilling holes into the outer casing of the printer to connect the 37°C incubator and humidified 5% CO_2_ (**Figure 1 A**). Then, the shape and size of the reservoir containing the unpolymerized bioink (or hydrogel) were adjusted. For this purpose, a suitable multi-well bioprinting vat with optical properties similar to the original vat was fabricated by a attaching a fluorinated ethylene propylene (FEP) foil to the sticky side of a bottomless eight well-plate (Ibidi “sticky slide”) (Figure 1 B). A platform designed to fit into the well-plate was 3D printed with an unmodified Anycubic Photon S and a standard SLA resin (Figure 1 C and CAD file provided in the Supporting Information **Figure S1**). Aluminum cubes were glued on the platform so that the printed resin and glue would not be in contact with the bioink containing the cells, thereby reducing the risk of chemical toxicity for the cells. Next, a guide for correct placement of the constructs on the screen was designed, so that the projection on the LCD screen aligned with the newly designed platform (Figure 1 D and CAD file provided in Supporting Information Figure S1). To correctly positions the constructs on the slicing software, the guide was uploaded on the software, then the CAD files were placed in the squares of the guide, automatically aligning them with the platform. Finally, a custom plate holder was designed to fix the Ibidi eight well-plate within the printing chamber and align it with the pattern on the LCD screen (Supporting information, Figure S1). Any type of well plate could be used in combination with a customized platform. For example, we designed a printing platform suitable for a 96 well-plate (CAD file in Supporting Information, Figure S9). Similarly, the printing platform can be adapted to many other plate formats.

**Figure 1.**
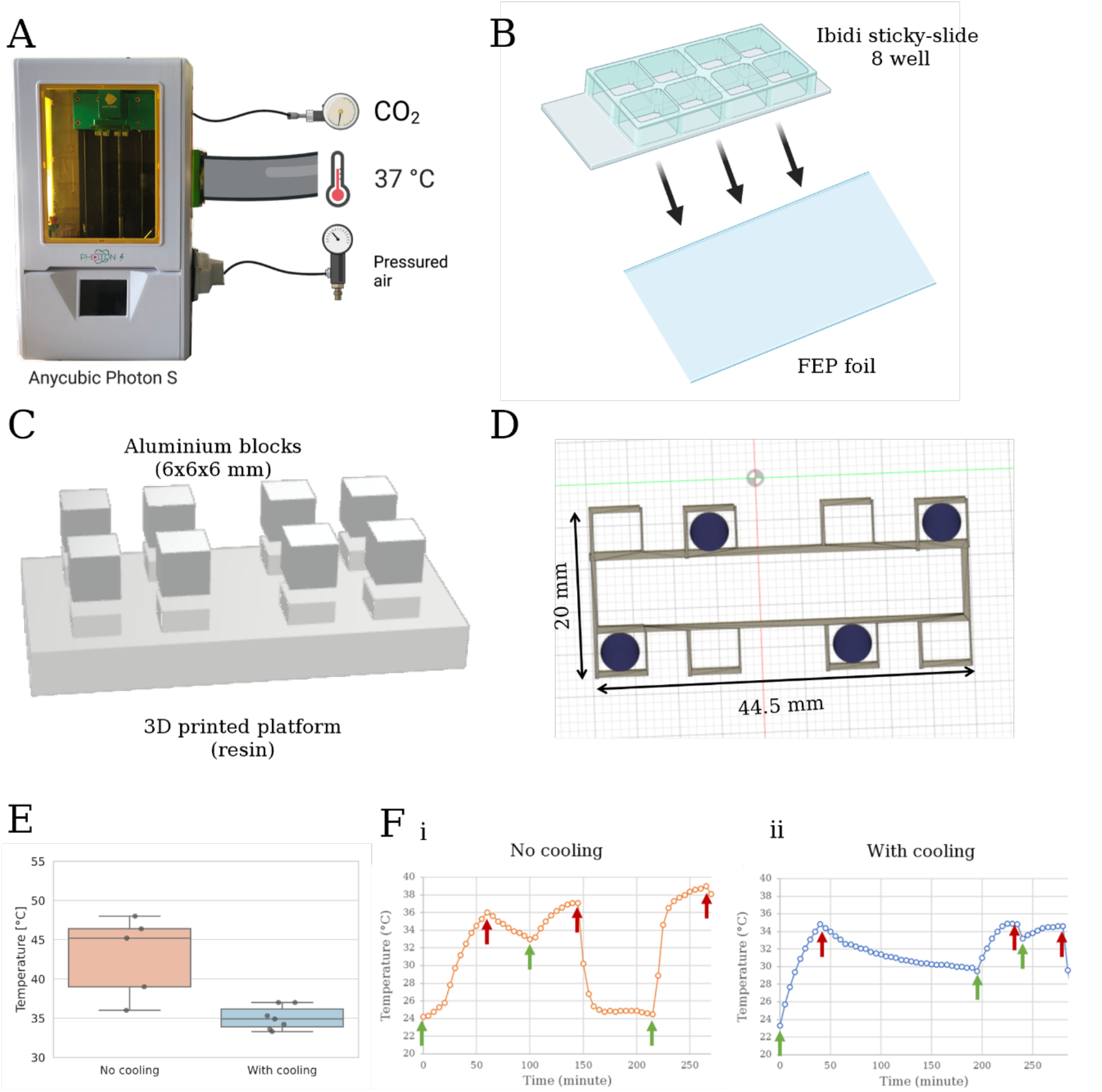
The Anycubic 3D printer is upgraded to a functional 3D bioprinter by integrating temperature control and CO_2_ incubation. A custom-made platform is designed to fit an eight well-plate to allow printing of several constructs in parallel. An air-cooling system maintains the LED screen at a constant temperature. (A) Illustration of the Anycubic Photon S printer with incubation and cooling modifications. Graphic created with BioRender.com. (B) Attachment of the Ibidi sticky-slide eight well-plate to a FEP foil provides identical optical properties to the material of the printer’s original vat bottom. Graphic created with BioRender.com. (C) CAD rendering of the 3D printed platform with eight aluminum plates for optimal hydrogel attachment and cell viability. (D) CAD rendering of the template (in grey) used to correctly align the constructs (in dark blue) with the platform. As an example, only half of the available eight slots were used to print cylinders. (E) The temperature on the surface of the uncooled LED screen reaches up to 48°C with a large variance (orange). Pressurized-air cooling (blue) keeps the temperature below 37°C and reduces the variance. (F) The temperature when printing consecutively is more stable with the addition of an air-cooling system. Green arrows indicate the start and red arrows the end of each print. (i) Without cooling, the temperature increases with each successive printing run. (ii) With the pressured-air-cooling, the temperature consistently remains below 37°C.

Preliminary tests showed that although the air temperature in the printing chamber was controlled at 37°C, the temperature at the surface of the LCD screen raised up to 48°C (Figure 1 E), increasing at each consecutive run (Figure 1 F i). An air-cooling system with pressurized air blowing directly onto the LED array was then added to the hardware (Figure 1 A). The cooling system ensured a precise control of the temperature at 37°C in the bioprinter (Figure E and F ii). A picture of the finished bioprinter can be found in Supporting Information **Figure S2**.

To encapsulate cells, the light dose applied to the system by the 3D bioprinter should be as low as possible while printing as fast as possible. The light dose of the screen when printing was measured during a typical run and found to be 52 mJ cm^-2^. An LED handheld lamp illuminating at 405 nm was used to encapsulate cells in a drop, thus showing the effect of near-UV radiation without the influence of the bioprinting process. The intensity of the light dose of the handheld lamp was also measured with the same photometer to be 414 mJ cm^-2^, about a factor of eight higher than the one in the bioprinter.

In this work, commercial devices were used to control temperature and CO_2_ in the printer. However, several tutorials to create DIY systems are available on the web (for instance open-source incubators^[35]^ at a cost of about $300, see https://amchagas.github.io/open-source-toolkit/post/diy_co2_incubator_bioreactor_for_mammalian_cell_culture__pelling_lab/). This allows to further decrease the cost of the set-up. The consumer SL printer is priced at around $300. The modifications elaborated in this work can be adapted to other SL printers (for example, Photon Mono SE or Mono X from Anycubic). The adapters to accommodate the multi-well plate are 3D-printed as well. Therefore, the bill of material has an overall cost well below €1,000.

## 3. The SL bioprinter permits the production of well-defined complex constructs

The manufacturer of the original 3D printer specifies a resolution of 1.25 µm in the x- and y-axes when using the SL resin. However, when using a new type of resin or hydrogel, it is advised to test the specifications necessary to obtain the desired resolution. In this work we used a photo-crosslinkable hydrogel composed by gelatin methacrylate (GelMA) and polyethylene glycol diacrylate (PEGDA) mixed at different concentrations. GelMA and PEGDA are often used in the literature due to the biocompatibility of the former and the structural properties of the latter.^[33,36–39]^ Several constructs suitable for bioengineering applications were printed as a proof of concept.

First, a lattice structure suitable for the use with sacrificial bioink to mimic blood vessels was printed. Sacrificial bioinks are degradable hydrogels (e.g., by heat, chemical or enzymatic digestion) that after being embedded in a second type of non-degradable hydrogel are dissolved, thus facilitating the fabrication of channels or vessels.^[40–45]^. The hydrogel used was 3%/3% GelMA/PEGDA. As shown in **Figure 2 A**, the grid features are well resolved, even though a thin layer of printed hydrogel was present across the bottom of the construct, due to over-polymerization of the bottom layer (the first three layers are over-exposed up to eight times more than the following layers to ensure adhesion to the platform). Next, a cube featuring a hollow channel was printed using a 10%/10% GelMA/PEGDA hydrogel (Figure 2 B). The assembly of hollow constructs is an alternative to sacrificial bioink to create blood vessels. The resolution of the cube was satisfactory with only the edges of the cube exhibiting slight rounding. The presence of a hollow channel, usually difficult to obtain by stereolithography, was very well defined in this case. Finally, structures mimicking microvilli lining intestinal epithelia were printed with a 10%/7% GelMA/PEGDA mixture (Figure 2 C). For this, over-polymerization on the bottom layers was exploited to obtain a base layer that was not present in the original design. Additionally, the over-polymerization allowed for the base of the pillars to be thicker than the top, further replicating the *in vivo* structure of a microvillus (Figure 2 C top row versus bottom row). The bioprinting process to obtain eight 60 mm^3^ constructs required approximately 30 minutes with the modified Photon S. The constructs printed were kept in PBS at 37°C for seven days and the structures neither collapsed nor decomposed. The constructs with high concentration of GelMA and PEGDA (10%/10% cube) required shorter light exposure time and therefore showed less over-polymerization than the ones with lower concentration (3%/3% lattice).

**Figure 2.**
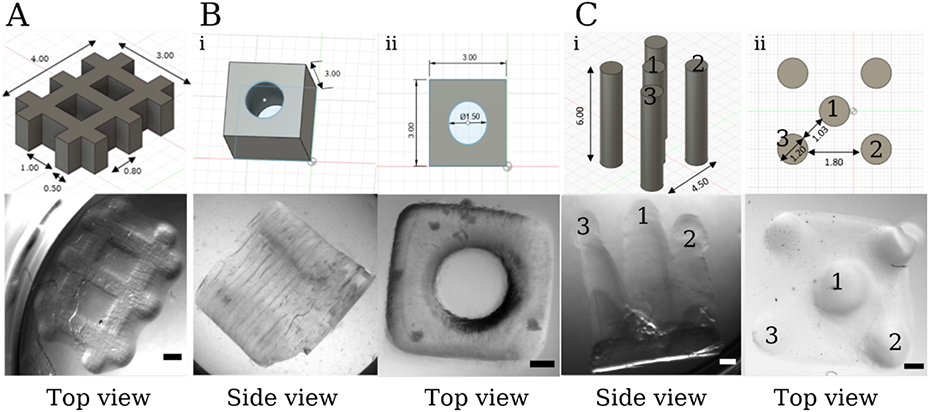
3D bioprinted constructs of variable shapes (lattice, hollowed, pillars) for different applications (sacrificial hydrogel, perfusable construct, microvilli). Top row: CAD design, bottom row: stereomicroscope images. (A) A 4 mm by 3 mm lattice is printed using 3%/3% GelMA/PEGDA hydrogel. The central gaps measuring 1 mm by 0.8 mm (top row) are well resolved except for a thin layer of over-polymerization at its bottom (bottom row). (B) A hollow cube is bioprinted using a 10%/10% GelMA/PEGDA hydrogel. The channel is entirely hollow and with a well-defined contour. (i) Side view of the cube showing the constant diameter of the hollow channel. (ii) Top view. (C) Structures resembling microvilli of intestinal epithelia. The typical microvillar structure is obtained taking advantage of the over-polymerization occurring on the bottom layers of the construct. (i) Side view distinguishing the two side (2 & 3) and central pillars (1). (ii) Top view highlighting the three pillars shown in (i). Scale bar: 500 µm. Microscope: Zeiss SteREO Discovery V8. Objective: Plan Apo S, 0.63x FWD 81 mm. Camera: AxioCam IcC SIN. Pixel size: 4.54 x 4.54 µm².

## 4. Characterization of the different hydrogel formulations guides the bioprinting process

The hydrogel used for 3D bioprinting was a mixture of GelMA and PEGDA at different ratios. This hydrogel is commonly used in 3D bioprinting and tissue engineering ^[38,46–48]^ since GelMA and PEGDA are easily synthetized in the laboratory or purchased. The required material properties of the printed hydrogel constructs depend on the application. The investigated hydrogel formulations were 3%/1.5%, 3%/3%, 5%/3% and 7%/3% (GelMA/PEGDA). The PEGDA concentration was kept low (1.5% or 3%) to limit the amount of synthetic hydrogel that is not degradable by the cells. To characterize those various formulations, we tested the hydrogels rheology, refractive index, and swelling properties.

First, to check for possible optical aberrations arising during the printing process, the refractive index of the different hydrogel formulations was measured (**Figure 3 A**). As expected, the refractive index of the hydrogels increased with increasing concentration of the components. However, the values remained very close to the refractive index of FEP foil (1.341–1.347), the material of the well-plate’s bottom. Therefore, the refractive index mismatch during the printing process was negligible for all practical purposes.

**Figure 3.**
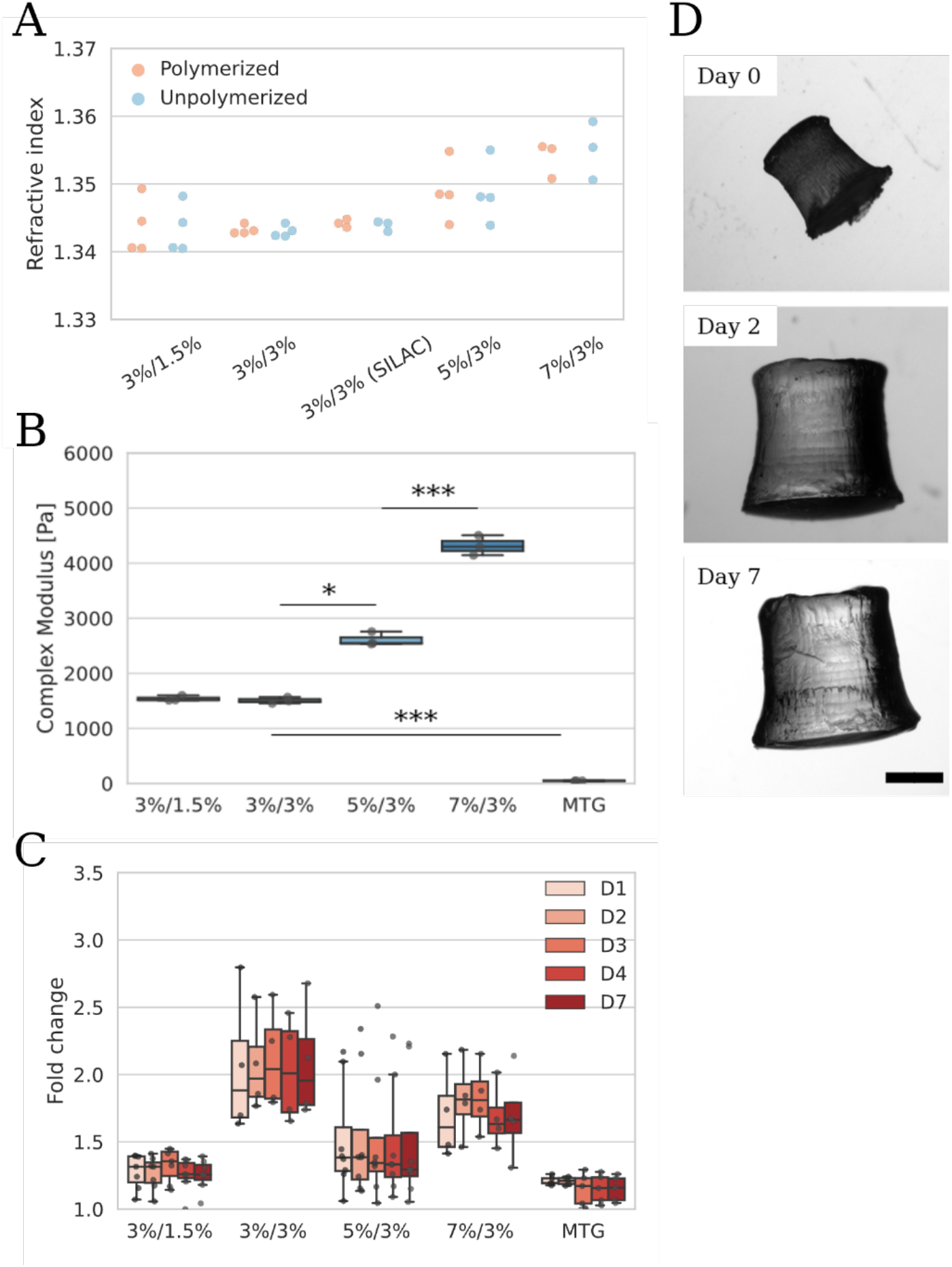
Rheological characterization of the various hydrogel formulations is used to determine the optimal hydrogel ratio for a given application. (A) The refractive index of the hydrogels is close to that of FEP foil bottom of the printing containers, thus minimizing refractive index mismatch. (B) The hydrogel’s complex modulus is adjusted by changing the GelMA concentration. PEGDA did not significantly influence the complex modulus. Interestingly, the modulus of 3%/3% GelMA/PEGDA differs significantly from that of Matrigel. (C) The swelling ratios of the hydrogels does not significantly vary. The values were normalized to day zero. The hydrogel formulation with the highest fold change in weight is 3%/3% GelMA/PEGDA, which corresponds to a higher retention of water in the crosslinked meshwork. (D) Pictures of 3%/3% GelMA/PEGDA cylindrical constructs printed for the swelling testing, on day zero, day two and day seven, illustrating the high amount of swelling after 48 hours, followed by a relative stability. Scale bar: 1000 µm. Microscope: Zeiss SteREO Discovery V8. Objective: Plan Apo S, 0.63x FWD 81 mm. Camera: AxioCam IcC SIN. Pixel size: 4.54 x 4.54 µm². The data is tested for significance with Student’s t test, n=3, p<0.005 (***), p<0.01 (**) and p<0.05 (*). Plot reading key: first the GelMA concentration is written, then the PEGDA concentration (for example, 3%/1.5% referred to 3% GelMA and 1.5% PEGDA). 3D bioprinted objects are referred to as Print. Matrigel is referred to as MTG.

Next, the shear complex modulus of the hydrogels was measured using a rheometer. The complex (or dynamic) modulus contains the elastic (G’) and the viscous (G”) modulus.^[49]^ It defines the main properties of a hydrogel, namely the capacity to absorb and dissipate energy. Measuring the complex modulus is fundamental in biomaterial engineering to best replicate the mechanical properties of the tissue of interest. The complex modulus of the GelMA and PEGDA hydrogel was mainly determined by the GelMA content, since the complex modulus increased in a nearly linear fashion at increasing GelMA concentration, whereas at a constant GelMA concentration and at decreasing PEGDA concentration the complex modulus remained nearly constant (Figure 3 B). The complex modulus closest to a healthy liver was given by the 3%/3% GelMA/PEGDA formulation, while constructs closer to a fibrotic liver were provided by 7%/3% GelMA/PEGDA mixture.^[50][51]^ The constructs, no matter the concentration, also showed a higher elastic modulus than viscous modulus (G’>G”) which indicated high levels of cross-linking within the constructs^[52]^ (see Supporting Information, **Figure S3**). Matrigel, a murine-derived extracellular matrix extract routinely used for 3D cell culture^[53–56]^ was tested as a control. The complex modulus of the Matrigel sample was measured at approximately 50 Pa, consistently with previous studies using oscillatory rheometers.^[57, 58]^

Finally, the swelling properties of the bioprinted hydrogel were assessed (Figure 3 C & D). The swelling ratio of the hydrogel is an important parameter for 3D bioprinting. Swelling depends on the degree of crosslinking and the porosity of the hydrogel and influences both the bulk geometry and the molecular diffusivity in the construct.^[59]^ The swelling of the printed constructs was stable after 24 hours in PBS and at 37°C. The swelling reached 2 to 2.5 folds for 3%/3% GelMA/PEGDA, while the other formulations displayed a lower swelling.

The various hydrogel formulations provide a broad range of mechanical properties, proving to be highly versatile to mimic different tissue types. Indeed, tissues display a wide range of rheological behaviors (reflected by the complex modulus) and water retention (reflected by the swelling properties) that need to be replicated in physiologically relevant engineered constructs. For example, the healthy liver is a spongious organ with an elastic modulus between 0.64 and 1.08 kPa^[60]^, while the cholangiocarcinoma tumor’s is between 3 and 12.1 kPa.

## 5. Imaging of the hydrogels using cryo-focused ion beam scanning electron microscopy allows precise characterization of their structure and porosity

Hydrogel porosity plays a major role for the diffusion of nutrients and small molecules, as well as cell adhesion and migration.^[54–56]^ The characterization of the hydrogel’s porosity was thus essential to identify the optimal microenvironment for the cells. The GelMA/PEGDA samples as well as the Matrigel controls were imaged with a cryo-focused ion beam scanning electron microscope (cryo-FIB SEM). Cryo-FIB SEM provides better sample preparation and higher resolution imaging than other electron microscopy techniques.^[61]^ The samples were first plunge-frozen to better conserve the hydrogel structure. The samples were then etched on the surfaced using a high current focused ion beam (FIB) to eliminate the ice-covered superficial layers (see Supporting Information **Figure S4**). Finally, a FIB parallel to the milling direction captured the pores minimizing the artifacts created by etching.

The SEM images of the hydrogels display an interconnected network of polymer fibers forming small cavities with various pore sizes and volumes (**Figure 4 A**). The 3%/1.5% GelMA/PEGDA hydrogel samples exhibit higher heterogeneity within the sample, having both larger and smaller pores present in the region of interest.

**Figure 4.**
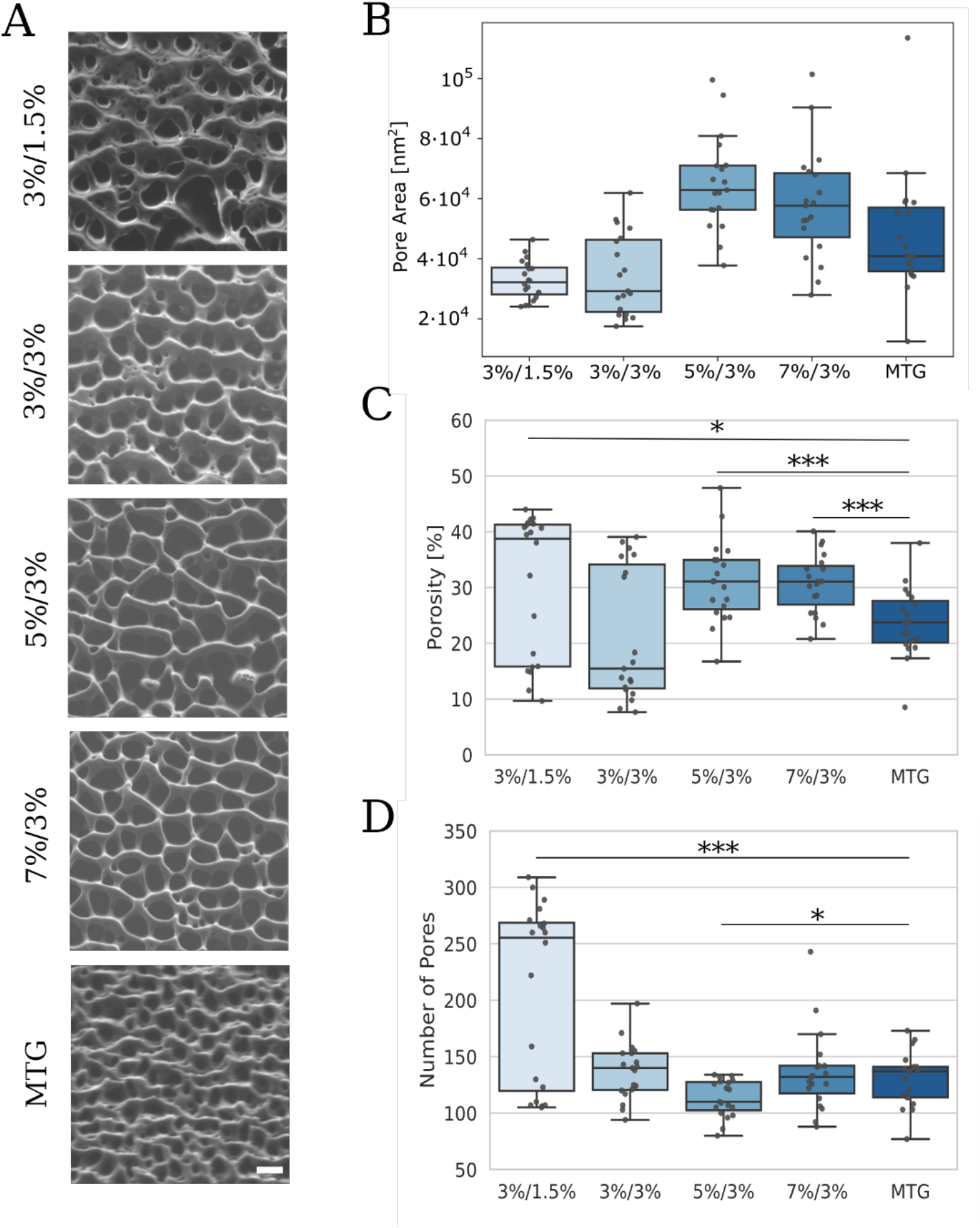
SEM characterization of the various hydrogel formulations compares pore properties (area, porosity and number of pores) of four hydrogel mixtures with that of Matrigel. (A) SEM images acquired by using cryo-FIB show interconnected fibers at all hydrogel concentrations and Matrigel. Scale bar: 500 nm. (B) The high-resolution cryo-FIB SEM images are analyzed by measuring the pore area with the DiameterJ plugin on ImageJ. (C) Quantification of the porosity (%). (D) Measurement of the number of pores. The data is tested for significance with a Student’s t test, 2 replicates per condition with n=10 for each replicate, p<0.005 (***), p<0.01 (**), p<0.05 (*). Plot reading key: first the GelMA concentration is written, then the PEGDA concentration (for example, 3%/1.5% referred to 3% GelMA and 1.5% PEGDA). 3D bioprinted objects are referred to as Print. Matrigel is referred to as MTG.

The median pore area of Matrigel was in the same range of the hydrogel samples, with no significant difference measured (Figure 4 B). The median porosity of Matrigel was significantly different from 5%/3% as well as 7%/3% (Student’s test, p<0.005) (Figure 4 C). The 3%/1.5% as well as 3%/3% hydrogels showed high heterogeneity between the samples with a broader range of porosity. Finally, the median number of pores in Matrigel was quite comparable to those of the hydrogel samples, except for 3%/1.5% which had significantly more pores (Student’s test, p<0.005) with a high heterogeneity between the different replicates (Figure 4 D).

## 6. CCAO cultured in hydrogel maintain high cell viability and their growth rate depends on hydrogel formulation

The viability of the CCA organoids as well as their morphology and growth in the different hydrogels were tested first without 3D bioprinting. The organoids were passaged and encapsulated in hydrogel drops with different formulation. Then, the drops were polymerized using a hand-held lamp with the same wavelength as the printer (405 nm) but with higher intensity, acting as a positive control for intense near-UV illumination conditions. The drops were then monitored for seven days with bright field microscopy. At day seven, the cell viability was measured with propidium iodide (PI, dead cells) and fluorescein diacetate (FDA, live cells). The morphology of the organoids observed during live imaging varied depending on the hydrogel formulation (**Figure 5 A**). At a low GelMA and PEGDA concentration (3%/1.5% GelMA/PEGDA), the cells did form organoids, but were highly mobile and migrated in the hydrogel and on the bottom of the plate, spreading further than the bound of the organoid. The organoids in the 3%/3% GelMA/PEGDA hydrogel showed a phenotype like those cultured in Matrigel. However, the organoids also displayed the tendency to form a compact structure, instead of a hollow lumen (Supporting Information **Figure S5**). At higher concentrations of GelMA (5% and 7%) the organoids showed more compactness compared to the Matrigel control. Finally, a drop of Matrigel was illuminated with a 405 nm handheld lamp to control whether illumination had an impact on the morphology, viability, and growth of the organoids. The resulting organoids showed identical morphology to the Matrigel control. The cell viability was overall high, with minimal cell death present in the sample.

**Figure 5.**
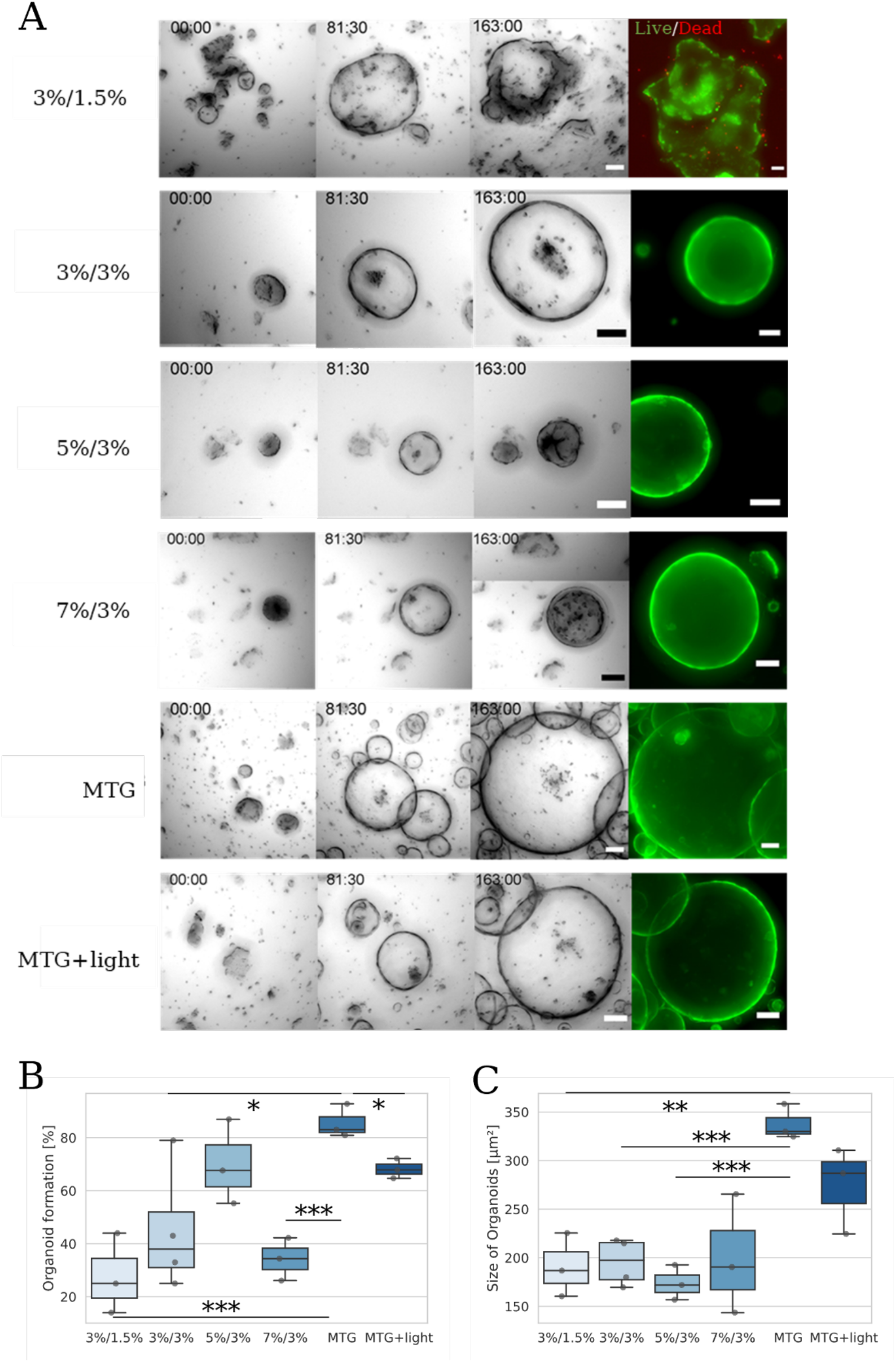
The morphology, viability and growth of the organoids cultured in different formulations of hydrogel varies from those of the organoids grown in Matrigel. (A) Brightfield images are acquired at different timepoints (0, 81:30 and 163 hours). After 168 hours, a live-dead assay is performed with PI (dead cells, in red) and FDA (live cells, in green). The organoids are viable in all the hydrogel drops. Scale bar: 100 µm. (B) The formation of organoids is calculated by counting the number of fragments on day zero and dividing this number by the number of formed organoids after seven days. The highest organoid formation rate (4-5%) is in Matrigel. (C) The size of the organoids is determined by randomly selecting eight to fifteen organoids per condition on day seven and averaging the values for each sample (conditions: n=3-4). The organoids cultured in hydrogel are smaller than the ones in Matrigel. Microscope: Zeiss Axio Observer Z1. Objective: Plan-Apochromat 5x/0.16. Camera: AxioCam MR R3. Voxel size: 1.29 x 1.29 x 60 µm³. Scale bar: 100 µm. The data is tested for significance with a Student’s t test, n=3 to 4, p<0.005 (***), p<0.01 (**), p<0.05 (*). Plot reading key: first the GelMA concentration is written, then the PEGDA concentration (for example, 3%/1.5% referred to 3% GelMA and 1.5% PEGDA). 3D bioprinted objects are referred to as Print. Matrigel is referred to as MTG. Matrigel samples that are exposed to 405 nm illumination as a control for near-UV potential damage unrelated to bioprinting are referred to as MTG+light.

Next, the growth rate of the organoids in the different hydrogel formulations as well as in the controls (Matrigel and the Matrigel exposed to 405 nm light) (Figure 5 B) was measured. The rate of the organoid formation was significantly different with respect to Matrigel in the 3%/1.5% as well as 7%/3% GelMA/PEGDA formulations (Student’s t-test, p<0.005). The organoids cultured in 3%/3% GelMA/PEGDA as well as the exposed Matrigel were also growing significantly less than in the Matrigel control (Student’s t-test, p<0.05). There was no significant difference in growth rate between 5%/3% GelMA/PEGDA and the Matrigel.

Finally, the size of randomly selected organoids was measured and compared to the Matrigel control (Figure 5 C). In all specimens the organoids were significantly smaller than in the Matrigel control except for the 7%/3% GelMA/PEGDA samples (Student’s t-test, p<0.01). The size did not vary significantly between the controls (Matrigel and the Matrigel exposed to 405 nm light).

Considering the results of the rheological characterization, electron microscopy images and the drop controls, we concluded that the optimal formulation for the model of cholangiocarcinoma was 3%/3% GelMA/PEGDA. From this point on, the term “hydrogel” refers to this formulation.

## 7. CCAO bioprinted in GelMA/PEGDA hydrogel are equivalent to organoids grown in Matrigel

CCA organoid fragments were suspended in 3%/3% GelMA/PEGDA and subsequently bioprinted with the modified SL printer. As a proof of concept, we printed a cylinder measuring 3.75 mm in height and 4.5 mm in diameter (**Figure 6 A**). The organoid growth in the printed construct was apparent after seven days in culture. Part of the organoids showed the typical monolayer morphology in both the bioprinted specimens and in the Matrigel control (Figure 6 B i, iii). However, most of them displayed a compact multilayer morphology with a small lumen (Figure 6 B ii). The same organoid morphology was observed in Broutier et al.^[62]^ and described as “resembling the corresponding tumor-of-origin”.

**Figure 6.**
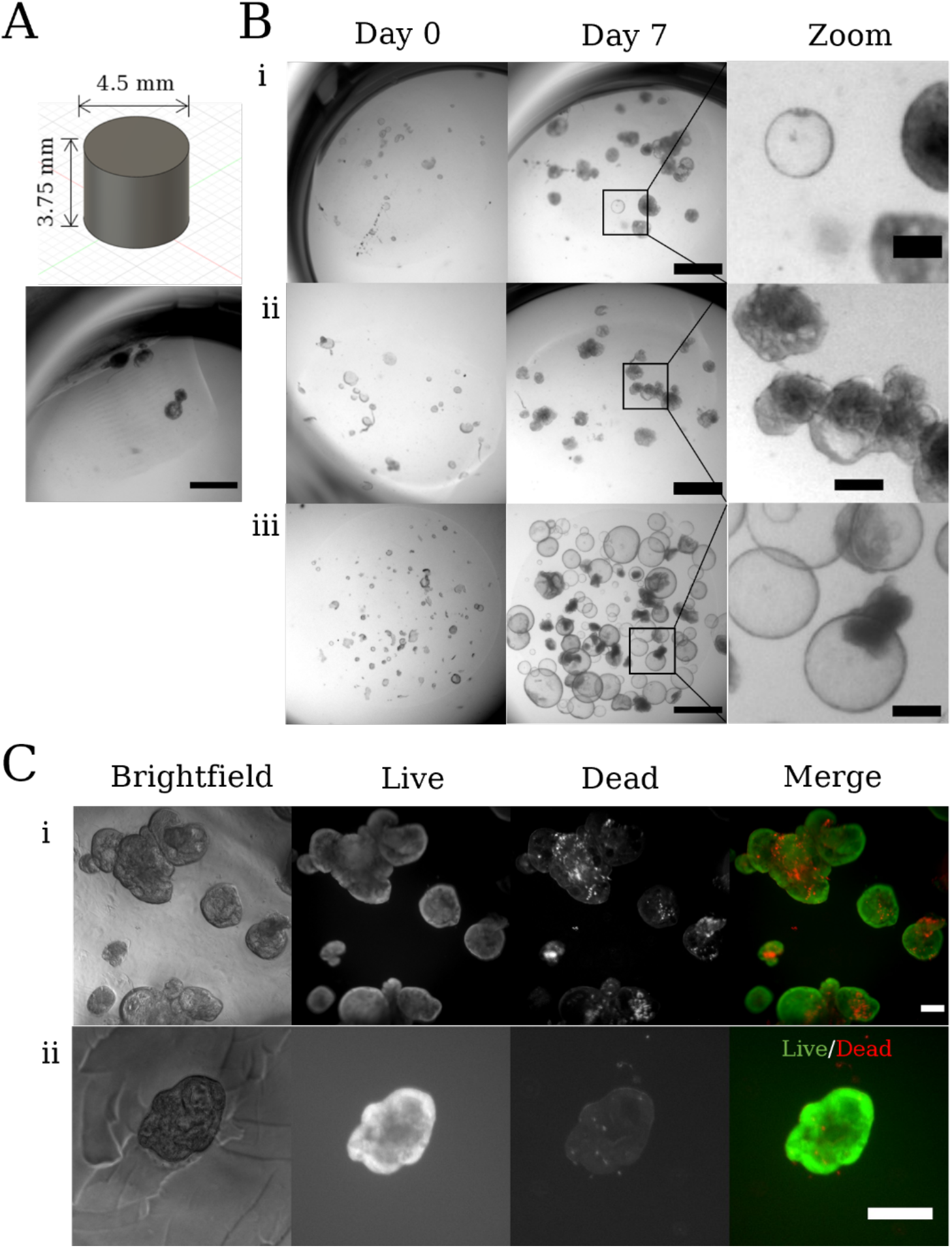
The morphology of the organoids cultured after 3D bioprinting mimics the one of the tumors of origin. The cells remain viable for at least seven days after printing. (A) CAD rendering of the 3D bioprinted constructs. The cylinder is a relatively complex shape that does not require a long printing time. (B) Organoid growth from day zero to day seven in two bioprinted constructs and in a Matrigel control. (i) Construct containing heterogenous organoids (scale bar: 1000 µm). A close-up of a monolayered organoid with hollow lumen is also shown (scale bar: 200 µm). (ii) Construct containing compact organoids (scale bar: 1000 µm). A close-up of an aggregate of compact organoids with smaller lumen is also shown (scale bar: 200 µm). (iii) Drop of Matrigel including monolayered organoids (scale bar: 1000 µm). A close-up of the structure of the organoids as well as a collapsed organoid is shown (dark group of cells) (scale bar: 200 µm). (C) Live-dead assay indicating the cell viability of the organoids seven days after bioprinting. Live cells are stained in green and dead cells in red (scale bar: 200 μm). (i) Aggregate of compact organoids with multiple smaller lumens shows cell death particularly at the organoid’s core. (ii) A single, smaller compact organoid displaying few dead cells. Microscope: Zeiss SteREO Discovery V8. Objective: Plan Apo S, 0.63x FWD 81 mm. Camera: AxioCam IcC SIN. Pixel size: 4.54 x 4.54 µm².

The viability of the organoids embedded in the bioprinted was tested on day seven after bioprinting (Figure 6 C). Although large aggregates (Figure 6C i) displayed some cell death in the central core, most of the cells were alive after seven days. Notably, smaller organoids (Figure 6C ii) exhibited a cell viability close to 100%. On average, the printed organoids showed 97% cell viability (Supporting Information **Figure S6**).

## 8. RT-qPCR analysis reveals no inflammation and limited DNA damage/oxidation in the bioprinted constructs

UV radiation is known to cause DNA damage such as double strand breaks. Moreover, the light exposure during bioprinting could lead to mutations in CCAO. Near-UV radiation (405 nm) has been shown to induce less DNA damage.^[63]^ However, the amount of DNA damage and additional stress markers present in the cells is highly dependent on the dose applied during bioprinting.^[64]^ To verify this, a RT-qPCR panel comprising 92 genes of interest and 3 reference genes, (glyceraldehyde 3-phosphate dehydrogenase - *GAPDH*), Hypoxanthine Phosphoribosyltransferase 1 - *HPRT1*, and Glucuronidase Beta -*GUSBI*) was used to evaluate the activity of DNA repair mechanisms. We screened 3D printed CCAO in a 3%/3% GelMA/PEGDA hydrogel and light-exposed Matrigel. The results were normalized with the gene expression measured in the Matrigel control (**Figure 7 A**).

**Figure 7.**
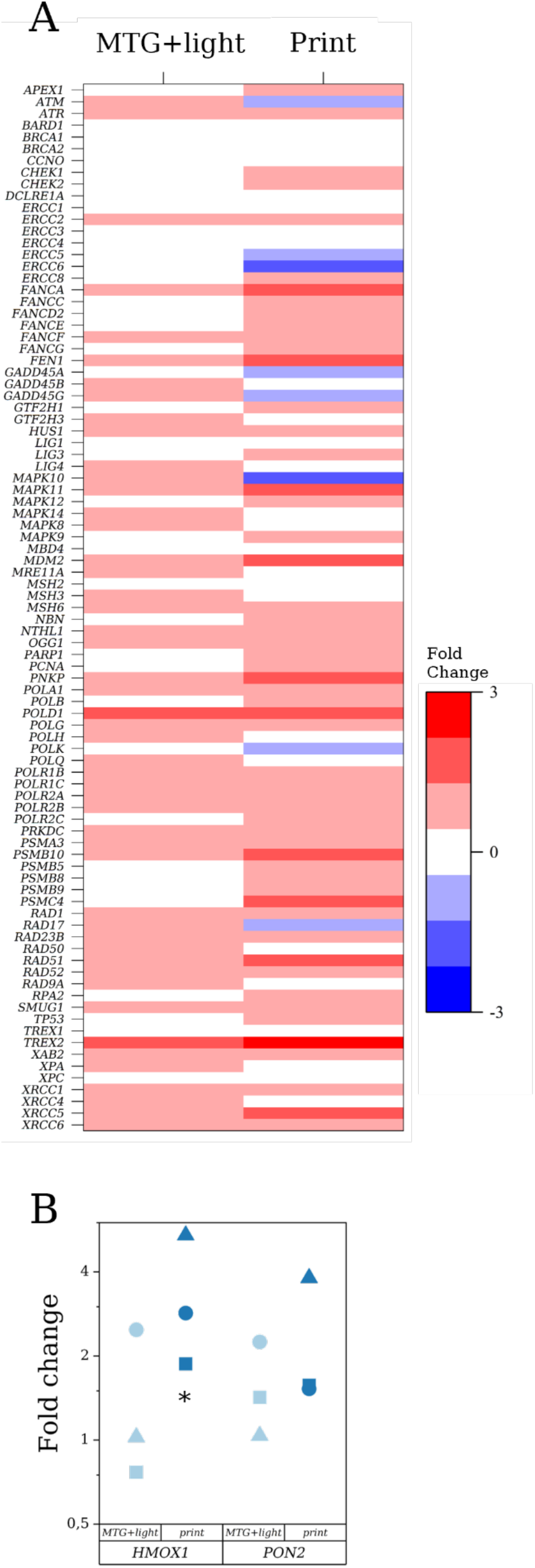
A RT-qPCR screening of DNA damage and reactive oxidative species markers indicates that stress is limited and not induced by near-UV illumination. (A) Heat map of RT-qPCR of 89 genes involved in DNA damage for light-exposed Matrigel (MTG+light) and the printed constructs (Print). The results are normalized to the Matrigel samples. The fold changes represent the level of up- or downregulation compared to the expression in the control sample (n=4 to 16). (B) RT-qPCR of two genes (HMOX-1 and PON2) involved in the response against reactive oxidative species. The samples are normalized to the Matrigel samples. HMOX1 is upregulated while PON2 only shows upregulation for one sample out of three. Each biological replicate (n=3 to 4) is a mix of 4 to 8 technical replicates. The data is tested for significance with a Student’s t test, p<0.05 (*). Plot reading key: print refers to the 3D bioprinted objects, MTG+light to the exposed Matrigel control.

Of the 92 examined genes, three could not be evaluated and only two were upregulated by a factor of >2. No gene was downregulated by a factor higher than 1.6 in the CCAO embedded in the printed hydrogel constructs. The difference between the CCAO in light-exposed Matrigel compared to not-exposed Matrigel was negligible. This suggests that the near-UV irradiation was not involved in the low-level DNA damage and that another factor in the bioprinting process (for instance residual unpolymerized monomers) led to the activation of DNA repair mechanisms.

As the photopolymerization of the hydrogel involves the formation of free radicals, the expression of the reactive oxygen species (ROS) markers heme oxygenase-1 (*HMOX1*) and paroxonase-2 (*PON2*) was investigated (Figure 9 B). A significant upregulation in the printed hydrogel constructs could be observed compared to both exposed and non-exposed Matrigel. However, only one sample (circle) showed an upregulation of PON2 in the exposed Matrigel compared to the not-exposed Matrigel. Interestingly, no differential expression of Interleukins 1A, 6 and 8 was detected (see Supporting Information **Figure S7**). Taken together, these results indicate that some level of oxidative stress and no inflammation were activated by the bioprinting.

We verified that photo-induced cell damage was limited in our system, as indicated by the negligible changes in the expression of genes involved in inflammation and oxidative stress between exposed and not-exposed Matrigel. However, the putative presence of free radicals in the unpolymerized hydrogel had an influence, as indicated by upregulation of HMOX1. Since two bioprinting runs were performed with one single batch of hydrogel/cell mixture (to save preparation time and material), a deterioration of the hydrogel components could explain the detected oxidative stress in the printed constructs compared to the illuminated drop.

## 9. CCAO cultured in the 3D bioprinted constructs maintain self-renewal and stemness characteristics

3D bioprinted tissues aiming to replicate the tumor environment should retain tumor-like characteristics of self-renewal and stemness. Typical tumor characteristics include high proliferation and low cell death presence as well as stem-cell markers.^[65]^

Ki67 was used as a marker for tumor proliferation^[66]^ and cleaved caspase 3 (Casp3) as a marker for apoptosis.^[67]^ High proliferation (and by association low apoptosis) levels can be used as a predictive tool for the assessment of tumor biopsies from patients.^[66]^ The cells in the printed constructs presented some proliferation, with levels similar to the controls (**Figure 8 A**). In the Matrigel controls, Ki67 was homogenously distributed throughout the organoids. In the bioprinted constructs Ki67 was localized at the tip of the organoid’s buddings, showing collectively coordinated growth. The ratio of proliferating cells to the total amount of cells was close in both printed and Matrigel samples. The ratio was slightly decreased in light-exposed Matrigel and print (Figure 8 B). We also observed a significant downregulation of the proliferation marker *MKI67* between the printed construct and the Matrigel control (Student’s t-test, p<0.005). In contrast, the exposed Matrigel and the Matrigel control showed similar expression levels (Figure 8 C). It has been shown that levels of mRNA were more sensitive as a prognostic tool than image analysis, in which case the RT-qPCR results should be the tool of choice in prognostic assessment.^[68]^ The staining against cleaved Caspase 3 (Casp 3) showed no apoptosis in any of the tested samples (Figure 8 D).

**Figure 8.**
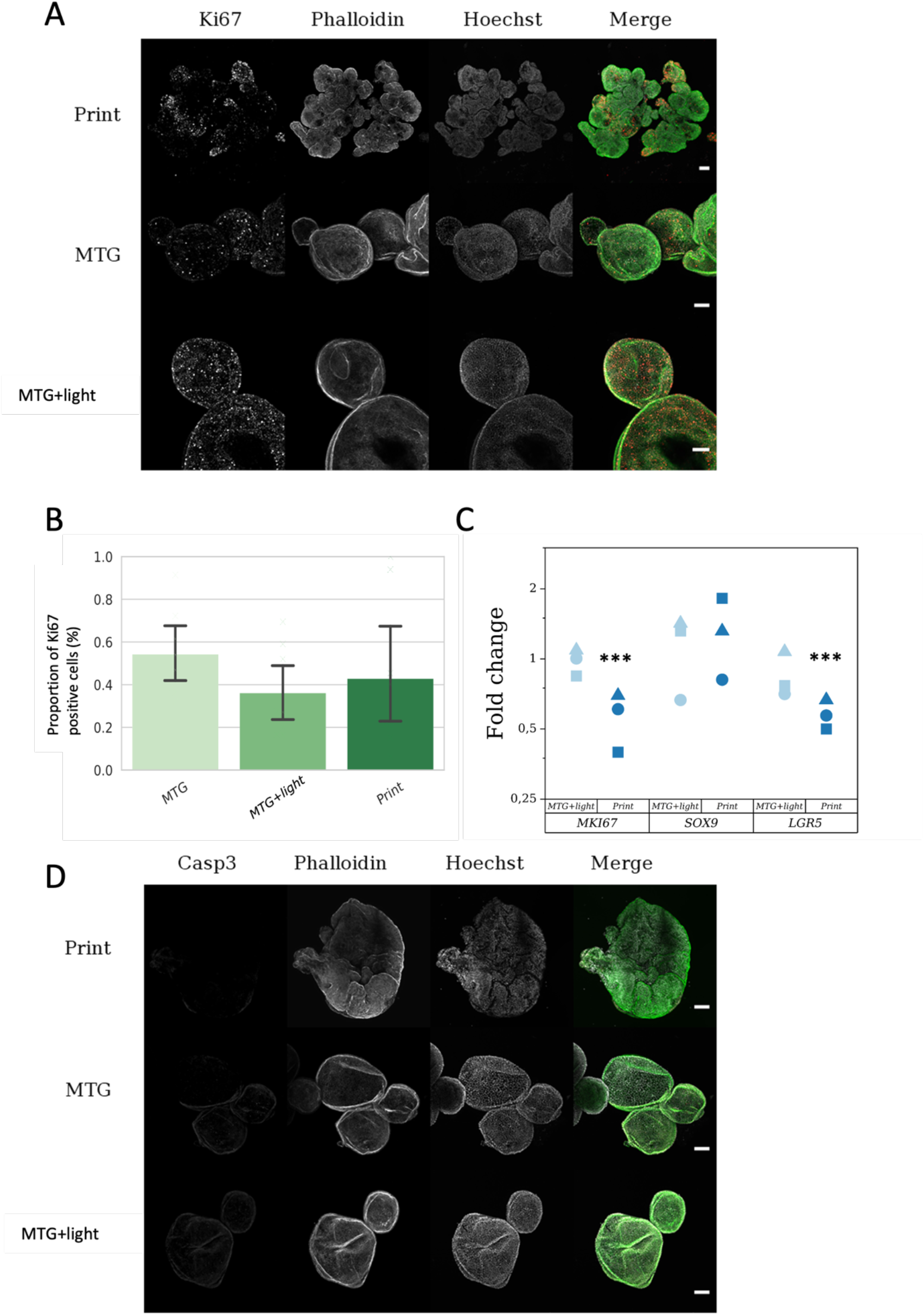
Presence and quantification of tumor self-renewal and stemness markers in the bioprinted constructs. (A) Ki67 localizes at the tips of budding organoids in the printed constructs whereas it is more diffuse in Matrigel and exposed Matrigel. Images of representative constructs stained against Ki67 (λ_ex_: 488 nm, λ_em_: 490-562 nm), Phalloidin for the cytoskeleton (λ_ex_: 561 nm, λ_em_: 568-644 nm) and Hoechst 33342 for the nuclei (λ_ex_: 405 nm, λ_em_: 410-502 nm). (B) Quantification of the ratio of proliferating cells (cells expressing Ki67) to the total amount of cells (stained with Hoechst 33342). (C) RT-qPCR analysis of the expression of MKI67 (proliferation marker), SOX9 and LGR5 (stem-like markers). MKI67 is downregulated in print compared to Matrigel. SOX9 and LGR5 are expressed and up- and downregulated, respectively. The samples are normalized with respect to the Matrigel samples. The data is tested for significance with a Student’s t test, n=3, p<0.005 (***), p<0.01 (**), p<0.05 (*). (D) No apoptosis is detected in the printed construct or the Matrigel and exposed Matrigel samples. Images of representative constructs stained against Caspase 3 or Casp3 (λ_ex_: 561 nm, λ_em_: 568-633 nm), Phalloidin for the cytoskeleton (λ_ex_: 633 nm, λ_em_: 697 nm) and Hoechst 33342 for the nuclei (λ_ex_: 405 nm, λ_em_: 410-502 nm). Microscope: Zeiss AxioObserver LSM780. Objective: Plan ApoChromat 20x/0.8 M27. Voxel size: 0.92 x 0.92 x 4 µm. Scale bar: 100 μm. Number of samples: n=2 to 3. 3D bioprinted objects are referred to as Print. Matrigel is referred to as MTG. Matrigel samples that are submitted to 405 nm illumination as a control for near-UV potential damage are referred to as MTG+light.

Stemness is crucial for drug resistance and relapse in many cancer types.^[69]^ Cholangiocarcinoma cells express the stemness markers Sex-determining region Y-box (SRY-box) containing gene 9 (*SOX9*)^[70]^ and leucine-rich repeat-containing G protein-coupled receptor 5 (*LGR5*).^[71]^ The expression of these markers was quantified with RT-qPCR (Figure 8 C). The marker *SOX9* was expressed similarly in the printed construct and the Matrigel controls as well as the exposed Matrigel samples. *LGR5* was significantly downregulated in the printed constructs when compared to the Matrigel controls, unlike the exposed Matrigel which showed similar expression levels.

Taken together, these results show that the organoids in the bioprinted construct maintain self-renewal characteristics, as shown by the presence of proliferation and stemness markers. Downregulation of LGR5 and equal expression of SOX9 compared to Matrigel are consistent with RNAseq measurements made by Broutier et al. 2017^[62]^ (comparison of the original tumor from the patient and the organoids cultured in Matrigel). These results suggest that the bioprinted constructs are accurately mimic of the original tumor biopsy.

## 10. Cancer prognostic and mechano-sensitivity markers are detected in the bioprinted constructs

Cholangiocarcinoma is known to be positive for Cytokeratin-19 (*KRT19*, an epithelial marker) whereas its high expression generally correlates with a worse prognosis in patients.^[72]^ Therefore, CCA organoids cultured in Matrigel, in exposed Matrigel and in 3D printed 3%/3% GelMA/PEGDA hydrogel constructs were stained by immuno-fluorescence against KRT19. All three samples showed a cytoplasmatic KRT19 expression as confirmed by nuclei counterstaining with Hoechst (**Figure 9 A**). To validate these findings, RT-qPCR was performed (Figure 9 C). A significant increase in *KRT19* expression was found in bioprinted constructs compared to Matrigel (Student’s t-test, p<0.005) and a decrease in exposed compared to not-exposed Matrigel (Student’s t-test, p<0.05). Cytokeratin-7 (*KRT7*), associated with a poor diagnostic in cholangiocarcinoma patients^[73]^, was also investigated. The expression of *KRT7* was significantly increased in the bioprinted constructs compared to Matrigel (Student’s test, p<0.005).

**Figure 9.**
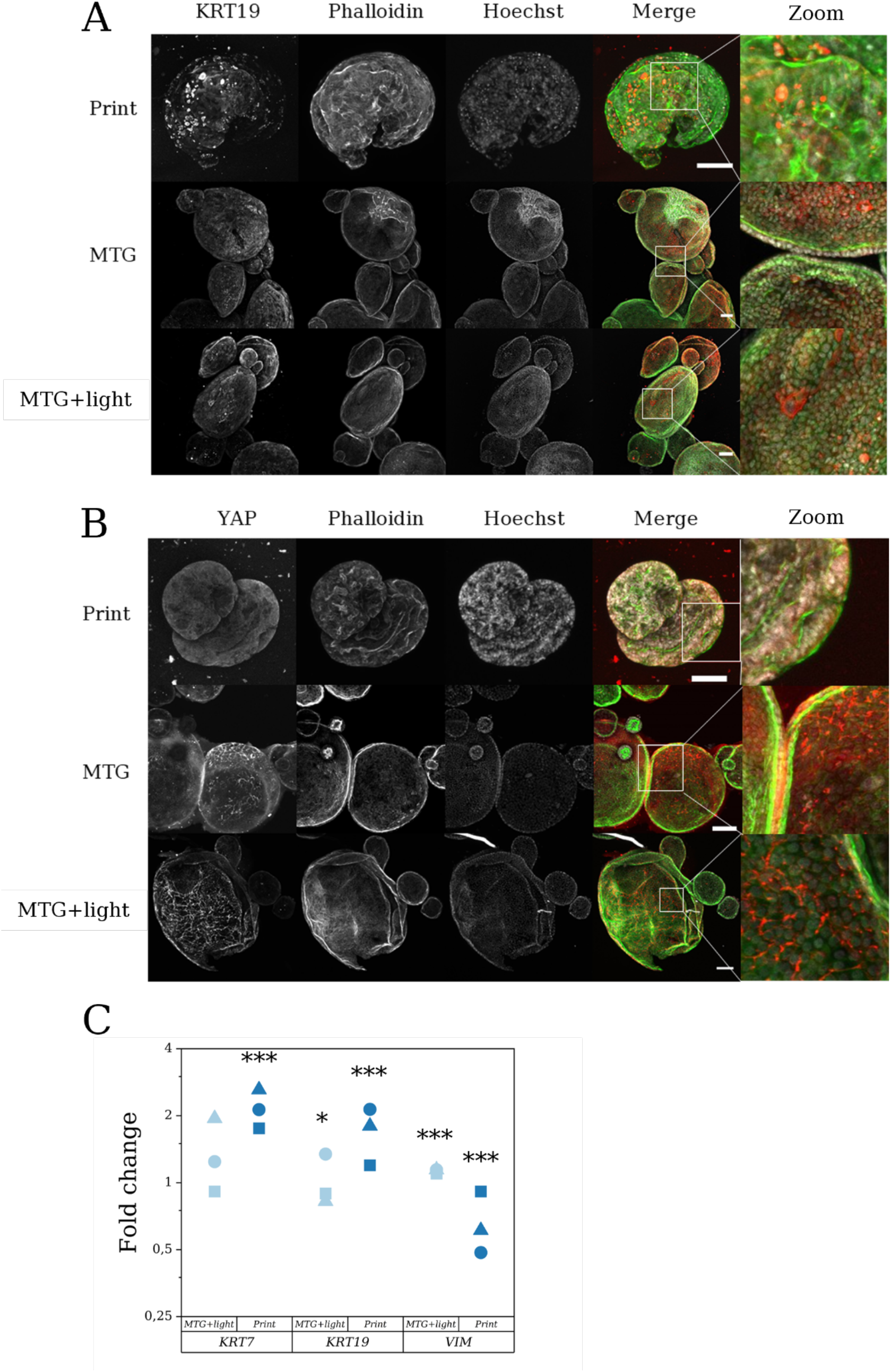
Detection of cancer prognosis (Keratin 19, Keratin 7 and Vimentin) and mechano-sensitive markers (YAP1) in the organoids. (A) The localization of Keratin 19 (KRT19) does not vary depending on the culture conditions. Images of representative constructs stained against Keratin 19 or KRT19 (λex: 488 nm, λem: 490-562 nm), Phalloidin for the cytoskeleton (λex: 561 nm, λem: 568-644 nm) and Hoechst 33342 for the nuclei (λex: 405 nm, λem: 410-502 nm). (B) Localization of YAP 1 does not differ between the printed and the Matrigel samples. Confocal images of representative constructs stained against Yes-associated Protein 1 or YAP (λex: 488 nm, λem: 490-562 nm), Phalloidin for the cytoskeleton (λex: 561 nm, λem: 568-644 nm) and Hoechst 33342 for the nuclei (λex: 405 nm, λem: 410-502 nm). (C) RT-qPCR analysis of markers associated with poor prognostic. Keratin 7 (KRT7) and Keratin 19 (KRT7) are upregulated while Vimentin (VIM) is downregulated. The samples are normalized to the Matrigel samples. The dataias tested for significance with a Student’s t test, n=3 to 4, p<0.005 (***), p<0.01 (**), p<0.05 (*). Microscope: Zeiss AxioObserver LSM780. Objective: Plan ApoChromat 20x/0.8 M27. Scale bar: 100 μm. Voxel size: KRT19: 0.69 x 0.69 x 4 µm³. YAP: 0.92 x 0.92 x 4 µm³. Plot reading key: 3D bioprinted objects are referred to as Print. Matrigel is referred to as MTG. Matrigel samples that are exposed to 405 nm illumination as a control for near-UV potential damage unrelated to bioprinting are referred to as MTG+light.

In contrast, the mesenchymal marker *VIM* coding for the intermediate filament Vimentin showed a significant downregulation in printed constructs compared to Matrigel (Student’s t-test, p<0.005, Figure 8 C). High expression of vimentin in CCA cells has been shown to correlate with a poor prognosis in patients.^[74]^

Yes-associated protein 1 (YAP) is a mechano-sensitive transcription factor that acts as a heterodimer with transcriptional co-activator with PDZ-domain (TAZ). YAP and TAZ are activated by the formation of stress fibers and stiffening of the extra-cellular matrix (ECM).^[75]^ Comparing the CCA organoids in the printed hydrogel constructs with the organoids cultured in exposed or not exposed Matrigel, no change in localization could be detected (Figure 9 B). YAP localized in the cytoplasm with a slightly higher concentration at the lateral cell membranes (cell-cell contact) in Matrigel-cultured organoids. The similar localization of the mechano-sensitive marker YAP confirmed that the stiffness of the hydrogel was not high enough to change localization of the protein.

These results show that differentiation (*VIM*) and epithelial (*KRT19* and *KRT7*) tumor markers are consistently expressed in the bioprinted CCA constructs. Additional markers supporting those results were tested with RT-qPCR (Supporting Information Figure S7 and **Figure S8**). Light exposure does not influence either the protein expression and localization or the mRNA genes expression as shown by the study of exposed Matrigel.

Interestingly, some of these results differed with those described in Broutier et al.^[62]^ in relation to the same organoid line. It is possible that the original donor’s liver had different mechanical properties (not measured in Broutier et al. and thus unknown). This can change the expression levels of the considered markers. However, the mechanical properties of the bioprinted GelMA/PEG hydrogel can be fine-tuned to properly match the characteristics of the original tumor *in vivo*.

## 11. Conclusion

We successfully modified an affordable consumer SL printer for bioprinting and showed that cells seeded in the printed GelMA/PEGDA hydrogel are highly viable with low levels of DNA damage. We also verified with RT-qPCR and immunofluorescence that the bioprinted constructs mimic key aspects of cholangiocarcinoma. Given its low-cost, small footprint and the room for customization, including the integration of high-throughput adapters (for 96 or even 384 well plates), we hope that this DIY SL bioprinter will pave the way for the widespread adoption of bioprinting in basic and translational research.

Further additions to improve the bioprinted constructs include the integration several cell types such as fibroblasts and macrophages to better replicate the tumor micro-environment. Experiments involving a similar tumor microenvironment including human breast cancer cells (T47D) and human dermal microvascular endothelial cells (HDMEC) in a 10%/10% GelMA/PEGDA hydrogel were performed and show promising results. The addition of a perfusion system could take advantage of the possibilities created by printing hollow channels. The addition of an XY-stage to translate the multi-well plate across multiple positions could be integrated with the bioprinter to allow the sequential layer-by-layer printing of multiple bioinks with different stiffness or containing other cell types.

## 12. Experimental Section/Methods

### Modification of a commercial SL 3D printer into a 3D bioprinter

An Anycubic Photon S printer (Anycubic Technology Co. Limited, Tsim Sha Tsui, Hong Kong) was modified. First, temperature and CO_2_ controls were installed. To do so, openings were cut in the case connecting the printing chamber to the heater (Solent Scientific Ltd, Portsmouth, UK) and to the CO_2_ control device (Carl Zeiss GmbH, Oberkochen, Germany). Then, a custom-made adapter was 3D printed (see Supporting Information Figure S1 for the CAD design) and connected to an opening cut on the side of the printer at a position below the LCD screen to direct a flow of pressurized air to the 405 nm LED array. Finally, a platform and a plate holder were 3D printed to be used with the Ibidi multi-well plates (CAD designs are available in Supporting Information Figure S1). Aluminum plates were glued onto the modified printing platform to foster hydrogel attachment and prevent any contact of the cells with the cytotoxic resin.

To check the efficiency of the temperature control in the printing chamber, a temperature data logger (Tempo Disc Bluetooth Temperature, Humidity and Dew Point Sensor, BlueMaestro, London, UK) was placed on the LCD screen and the temperature was measured during 3D printing, with and without air-cooling of the LED array. Next, several print runs were carried out in succession during temperature logging.

### Multi-well plates

We used a Sticky Slide 8-well µ-slide (Ibidi GmbH, Graefelfing, Germany). On the sticky side of the plate, a 50 µm fluorinated ethylene propylene (FEP) foil (DuPont de Nemours International SA, Geneva, Switzerland) was applied. The platform on which the constructs are printed was custom-designed and 3D printed to fit the 8-well plate. Moreover, a template guide was developed to allow a precise alignment of the constructs with the eight platforms (files available in the Supporting Information).

### Preparation of the hydrogel

The hydrogel used for this study was a combination of gelatin methacrylate (GelMA) gel strength 300 g, Bloom, 80% degree of substitution (Sigma-Aldrich Chemie GmbH, Steinheim, Germany) and polyethylene glycol diacrylate (PEGDA), average Mn 4000 (Sigma-Aldrich Chemie GmbH, Steinheim, Germany). The hydrogel components were separately mixed with either phosphate buffer saline (PBS, Gibco, ThermoFisher Scientific, Waltham, MA, USA) when the hydrogel was used for material characterization purposes, or with SILAC advanced DMEM/F-12 Flex Media, with no additives and no phenol red (Gibco, ThermoFisher Scientific, Waltham, MA, USA) when the hydrogel was used to encapsulate living cells. 0.2% w/v Lithium-Phenyl-2,4,6-trimethylbenzoylphosphinat (LAP, Sigma-Aldrich Chemie GmbH, Steinheim, Germany) was added to the PEGDA mixture and the GelMA and PEGDA/LAP mixtures were dissolved separately at 65°C under shaking conditions for two hours. Subsequently, the dissolved GelMA and PEGDA were mixed to reach the desired concentration, 0.0025% w/v tartrazine (Sigma-Aldrich Chemie GmbH, Steinheim, Germany) was added, and the mixtures were left at 37°C for another hour. The concentrations used in this study were 3% w/v GelMA with 1.5% w/v PEGDA, 3% w/v GelMA with 3% w/v PEGDA, 5% w/v GelMA with 3% w/v PEGDA and 7% w/v GelMA with 3% w/v PEGDA.

## 3D bioprinting of the hydrogel samples

The constructs were all designed with Fusion 360 (Autodesk Inc., San Rafael, CA, USA) computer-aided design (CAD) software and sliced with Chitubox (Chitubox, Shenzhen, Guangdong, China). The constructs encapsulating live cells were cylinders, 3.75 mm in height and 4.5 mm in diameter. Further items were printed with the hydrogel to demonstrate the feasibility of printing more intricate designs: a 4 x 3 mm^2^ grid, a 3 x 3 x 3 mm^3^ cube with a 1.5 mm channel, a model of microvilli of 4.5 x 6 mm^2^ and a model of liver nodules of 4.2 x 4.5 x 6 mm^3^. The settings of the slicing software for printing were 150 seconds bottom illumination (3 layers), 80 seconds exposure time, 150 µm layer height, 3 mm lifting distance and 100 mm/min speed (bottom lift, lift and retract speed). Indeed, a slower stage lifting speed was shown to distribute the organoids fragments more homogenously within the construct.

### Rheology of the hydrogel

The different hydrogel formulations and Matrigel were tested to characterize their mechanical properties (n=3 to 4). The rheology of the samples was measured using a Gemini 150 advanced rheometer (Bohlin Instruments GmbH, Pforzheim, Germany). The hydrogel samples were prepared as previously described and 200 µL were placed on the measurement plate of the device, which was previously warmed to 37°C to prevent the hydrogel from hardening. The settings were the following: −0.1 N auto tension, 1 Hz frequency, constant strain of 1%. After 40 seconds, a Bluepoint LED Eco 405 nm hand-held lamp (Hoenle AG, Gräfelfing, Germany) placed right next to the device was turned on (2.93 W cm^-2^) to allow for full crosslinking of the hydrogel. The measurement was stopped after a total of 4 minutes of recording. For the Matrigel samples, the stage was cooled down to 4°C and 200 µL of the gel were placed on the stage. The settings for the device remained the same as for the hydrogels except for the strain which was set to 2%. After nearly 5 minutes, the temperature of the stage was set to 37°C. The measurement was stopped after a total of 40 minutes of recording. The complex modulus was extracted and processed on Excel (Microsoft Corp., Redmond, WA, USA). The last 15 data points were averaged to determine the complex modulus at full cross-linking.

### Determination of the hydrogel’s swelling property

Hydrogels with 3%/1.5%, 3%/3%, 5%/3%, 7%/3% GelMA/PEGDA (w/v) were prepared as previously described. Matrigel was thawed on ice for 20 min and 40 µl of liquid Matrigel were placed as a drop into a 40 µm cell strainer (Corning, Tewksbury, MA, USA), which was weighed beforehand. Matrigel drops were polymerized for 20 min at 37 °C and afterwards incubated in PBS (Gibco, ThermoFisher Scientific, Waltham, MA, USA) for 7 days at 37 °C. Cylindrical hydrogel constructs were printed as previously described. Bioprinted constructs and Matrigel were weighed immediately after printing and after polymerization, respectively. The weight was then measured on day one, two, three, four and seven and the ratio between the “dry” sample (right after polymerization) and “swollen” samples incubated in PBS for the respective duration was calculated. Day zero values were used for normalization in data analysis.

### Refractive index measurement

The refractive index of the hydrogels was measured using an automatic refractometer (J257, Rudolph Research Analytical, Hackettstown, NJ, USA). The hydrogels formulations were prepared as previously described and additionally 3%/3% GelMA/ PEGDA (w/v) in SILAC medium was prepared (n=3). 100 µL was pipetted onto the analysis window of the refractometer. After measuring, the sample was polymerized using the curing 405 nm hand-held lamp (15 seconds) and the refractive index was measured again.

### Analysis of hydrogel pore architecture by cryo-FIB SEM

The carbon side of Quantifoil R1/4 Cu Electron Microscopy (EM) grids (200 mesh) (Electron Microscopy Sciences, Hatfield, PA, USA) was glow-discharged for 20 seconds. Within 10 minutes after glow discharging the EM grids were used for plunge freezing without any further processing. A hydrogel solution with a volume of 1 µL was added to the EM grid. The hydrogel was then polymerized by exposition to a 405 nm hand-held lamp (Comgrow, Shenzhen, China). Matrigel (BD, Heidelberg, Germany) was polymerized at 37°C. The samples were plunge-frozen into liquid ethane with the Vitrobot Mark IV (ThermoFisher Scientific, Waltham, MA, USA). The Vitrobot chamber was equilibrated at 100% rH and 20 °C before the start of the experiment.

The plunge-frozen EM grids were clipped into cryo-FIB autogrids (ThermoFisher Scientific, Waltham, MA, USA) and loaded into a pre-tilted TEM grid holder with shutter and additional cold trap (Leica) under liquid nitrogen using an EM VCT500 loading station (Leica Microsystems GmbH, Wetzlar, Germany). The sample holder was then taken up by the EM VCT500 manual transfer shuttle and, via a VCT dock (Leica Microsystems GmbH, Wetzlar, Germany), transferred into the Helios 600i Nanolab FIB/SEM dual beam instrument (ThermoFisher Scientific, Waltham, MA, USA) equipped with a band-cooled cryo stage (Leica Microsystems GmbH, Wetzlar, Germany) equilibrated at −157 °C. The EM grids were screened using SEM (3 kV, 0.69 nA) and FIB (gallium ion source, 30 kV, 7 pA). To sublime water ice out of the hydrogel the stage temperature was increased to −60 °C and left for equilibration for 60 minutes. Flat regions of the frozen hydrogel were chosen for etching at a net incident angle of 53° (55° stage tilt). Ten random areas, each 10 x 10 µm^2^ in size, were etched with a FIB current of 400 pA for 60 seconds to reveal the hydrogel pores. At the same angle images of the etched positions were acquired at 65 000x magnification with a FIB current of 7 pA afterwards (Secondary Electron Detection with an ETD).

The images were analyzed using the DiameterJ plugin on FIJI^[83]^, which has been designed to segment and analyze SEM images of fibrous objects (ImageJ version 1.53c, U. S. National Institutes of Health, Bethesda, Maryland, USA). The pore area (total number of black pixels counted in pores divided by the total number of pores in image), porosity (total number of black pixels divided by the total pixels in an image) and the number of pores of the hydrogels were determined. Two replicates per hydrogel condition were imaged, comprising each 10 pictures of random positions.

### Cholangiocarcinoma organoids culture

The cholangiocarcinoma organoid (CCAO) line was kindly provided by Monique M. A. Verstegen and Luc J. W. van der Laan (Department of Surgery, Erasmus MC Transplant Center – University Medical Center, Rotterdam, The Netherlands). The organoid culture protocol was adapted from Broutier *et al.*, 2017^[62]^. Briefly, the basal medium was composed of Advanced DMEM/F12 (Gibco, ThermoFisher Scientific, Waltham, MA, USA) with 1% penicillin/streptomycin (1%, Gibco, ThermoFisher Scientific Waltham, MA, USA), 1% Glutamax (Gibco, ThermoFisher Scientific Waltham, MA, USA) and 10 mM HEPES (Gibco, ThermoFisher Scientific Waltham, MA, USA). Growth medium consisted of basal medium supplemented with 1:50 B27 (Gibco, ThermoFisher Scientific Waltham, MA, USA), 1:100 N2 (Gibco, ThermoFisher Scientific Waltham, MA, USA), 1.25 mM N-acetyl-L-cysteine (MilliporeSigma, St. Louis, MO, USA), 30% Rspondin-1 conditioned medium (prepared from the culture of 293T-HA-RspoI-Fc cells obtained from Calvin Kuo, Lokey Center for Stem Cell Biology and Regenerative Medicine, Stanford University, CA, USA), 10 mM nicotinamide (MilliporeSigma, St. Louis, MO, USA), 10 nM (Leu15)-gastrin (MilliporeSigma, St. Louis, MO, USA), 50 ng mL^-1^ recombinant human EGF (PeproTech, Rocky Hill, NJ, USA), 100 ng mL^-1^ recombinant human FGF 10 (PeproTech, Rocky Hill, NJ, USA), 25 ng mL^-1^ recombinant human HGF (PeproTech, Rocky Hill, NJ, USA), 10 µM forskolin (Tocris Bioscience, Minneapolis, MN, USA) and 5 µM A 83-01 (Tocris Bioscience, Minneapolis, MN, USA). The organoids were passaged by disrupting the Matrigel with rapid up-and-down pipetting and centrifuging the Matrigel and cells in basal medium. The supernatant was removed, and the cells were resuspended in the required amount of Matrigel (BD, Heidelberg, Germany) – 20 µL per well for a 48 well-plate, 10 µL per well for a 96 well-plate. The Matrigel/cell suspension was seeded as a droplet at the bottom of a pre-warmed non treated well-plate. The Matrigel was polymerized at 37°C for around 15 minutes before incubating the drops in growth medium.

To prepare the cells for 3D bioprinting, they were removed from the Matrigel, fragmented, and centrifuged as previously described. The supernatant was discarded and the fragment pellet was mixed equally in the 3%/3% hydrogel and in Matrigel (controls), so that a passaging ratio of 1:8 was obtained. Two series of eight cylinders were printed out of the hydrogel/fragments mix. Half of the Matrigel controls were illuminated with the 405 nm handheld lamp (15 seconds for 20 µL drops, 10 seconds for a 96 well-plate through the plastic well plate). After printing, the constructs were washed in PBS containing normocin 1:500 (Invivogen, San Diego, CA, USA). The PBS was removed and the constructs as well as the Matrigel controls were cultured in growth medium supplemented with ROCK inhibitor (10 µM Y-27632, R&D systems, Rocky Hill, NJ, United States). After 24 hours, the medium was changed to a growth medium without ROCK inhibitor. Control drops were also prepared using the same process but seeded as a hand-pipetted drop in a well plate and polymerized with the 405 nm curing light.

### Time lapse, cell viability and immunofluorescence staining of the organoids

The cells were seeding as drops in either the different hydrogel formulations or in Matrigel or Matrigel exposed to the 405 nm hand-held lamp. The cells were imaged for seven days with an inverted fluorescent and brightfield microscope (Cell Observer SD, Carl Zeiss GmbH, Oberkochen, Germany). The dead cells were stained with propidium iodide (PI) and the live cells with fluorescein diacetate (FDA) (Sigma-Aldrich Chemie GmbH, Steinheim, Germany) and imaged with the same microscope. Quantification of the organoid growth was performed by counting the fragments on day zero and counting the formed organoids on day seven using the multipoint tool on the FIJI software. Three samples per condition were quantified. The organoid size was calculated by measuring the area in µm^2^ of 8 to 15 organoids per condition on day seven, then averaging the area for each biological replicate (n=3 to 4).

Printed constructs were imaged with a fluorescence stereomicroscope (SteREO Discovery V8, Carl Zeiss GmbH, Oberkochen, Germany). A live-dead assay was performed as previously described (n=6) and t the cell viability was quantified with FIJI/ImageJ by measuring the area of the live staining and the area of the dead staining before calculating the ratio of the dead area over total area.

For immunofluorescence, the organoids grown in Matrigel were fixed in 4% PFA (MilliporeSigma, St. Louis, MO, United States) in PBS for 30 minutes, then Cell Recovery Solution (Corning, NY, USA) replaced the fixative for one hour on ice. The organoids separated from the matrix were processed in parallel with the bioprinted constructs. The bioprinted constructs were fixed and stained according to an adapted protocol from Loessner *et al*., 2016^[76]^. Briefly, the samples were fixed in 4% PFA in PBS for 30 minutes. They were then washed three times in PBS and permeabilized in Triton X-100 (0.3% v/v) in PBS for 40 minutes at room temperature. After washing thrice with 0.1 M glycine (Carl Roth GmbH, Karlruhe, Germany) in PBS and thrice in PBS-T (0.1% Triton X-100 in PBS, MilliporeSigma, St. Louis, MO, United States) for 10 minutes each time, the samples were blocked for one hour in BSA (0.1%), Triton x-100 (0.2%), Tween-20 (0.05%) in PBS and incubated in the primary antibody solution overnight (see Supporting Information **Figure S10**). The next day, the samples were washed in 2% penicillin/streptomycin in PBS thrice for five minutes before incubating in the secondary antibody solution (see Supporting Information Figure S10) for two hours. After a final wash in PBS and penicillin/streptomycin, the cells were ready for imaging. Two biological replicates were imaged per condition, with at least 3 images per biological replicate.

Proliferating cells were stained against Ki67 and quantified by selecting three slices within the z-stack, at the beginning, in the middle, and at the end of the organoid. A gaussian blur filter was applied to each picture, a binary mask and a watershed was then used to obtain a clear segmentation (the threshold for the binary mask was selected for each image individually). The FIJI function Analyze Particles^[77]^ was used to count the objects. The results for each slice were averaged and the mean of the two biological samples were plotted.

### Quantitative reverse transcription PCR (RT-qPCR)

For RT-qPCR, total RNA was extracted from CCA organoids grown in Matrigel with and without UV illumination as control (n=4 per biological samples were pooled) using TRIzol (Invitrogen ThermoFisher Scientific, Waltham, MA, USA) and following the manufacturer’s instructions. The bioprinter constructs (n=15 to 16 per biological sample) were homogenized in TRIzol using a micro pestle in a 1.5 ml tube (Carl Roth GmbH, Karlruhe, Germany), following an adapted protocol from Köster *et al.* 2016.^[78]^ RNA concentration and purity was determined using the NanoPhotometer NP80 (Implen GmbH, München, Germany).

Maxima First Strand cDNA synthesis kit with dsDNase (ThermoFisher Scientific, Waltham, MA, USA) was used for synthesis of 5 µg of cDNA. cDNA was stored at −20°C until further usage.

For SYBR Green RT-RT-qPCR, the PowerTrack SYBR Green Mastermix (ThermoFisher Scientific, Waltham, MA, USA) was used. All the primers were designed on PrimerBlast^[79]^ (National Center for Biotechnology Information (NCBI), Bethesda, MD, USA) unless otherwise stated (see Supporting Information **Figure S11**) and ordered on Biomers.com (Biomers GmbH, Ulm, Germany). The primers were diluted to a final concentration of 4 µM and 10 ng of cDNA was used per 10 μl reaction. The RT-qPCR was carried out in triplicates. To test the expression of inflammation and oxidative stress markers, the Fast Advanced TaqMan MasterMix was used. The primers were purchased from ThermoFisher (ThermoFisher Scientific, Waltham, MA, USA). A total of 20 ng cDNA was used in 20 μl reaction volume, which was performed in triplicates. For analysis of human DNA repair mechanisms, a pre-set plate, TaqMan Array Human DNA Repair Mechanism (ThermoFisher Scientific, Waltham, MA, USA) was used in combination with the Fast Advanced MasterMix.

For analysis, the geometric mean of the triplicates was normalized to the geometric mean of the two reference genes *RPL13* and *TBP* as well as to CCA organoids grown in Matrigel. A list of all primers can be found in the Supporting Information Figure S11.

### Statistical analysis

All data and representative images presented were obtained from at least 2 independent experiments or biological samples, with multiple technical replicates. Statistics and quantitative RT-PCR plots were generated in OriginLab (version 2021b, OriginLab Corp., Northampton, MA, USA). Scientific plots were generated in Python 3.7 with the MatPlotLib^[80]^, Pandas^[81]^ and Seaborn^[82]^ libraries. For Boxplots the median and interquartile range is shown with lower whiskers ranging 0 – 25% and upper whiskers 75 – 100%. Tests for differences between two groups were realized with Student’s two-tailed unpaired t-test. No data points were excluded from the analyses. Significance was tested at p-values 0.05 (*), 0.01 (**) and 0.005 (***).

## Supporting Information

Supporting Information is available from the Wiley Online Library.

## Acknowledgements

LB designed the experiments, LH performed the swelling experiments, KNW and LB conducted the experiments using the CCA organoids. LH, KW prepared and imaged the cry-FIB SEM samples. FP designed the printer modifications. LB and FP wrote the manuscript with assistance from KNW and LH. AF supervised KW. EHKS provided support to research. FP conceived and supervised the research. All the authors read and revised the manuscript.

FP, EHKS, LB, KNW, LH, HW thank the EU Horizon2021 project BRIGHTER (Grant #828931) for funding. We thank the Frankfurt Center for Electron Microscopy (FCEM) and for their support in the SEM imaging of the hydrogel samples and Danica Hagen for the images of the constructs containing a channel. We also thank Cellendes GmbH, Helmut Wurst and Gabriele Di Napoli for their help in the rheological measurements of the hydrogels. We thank Monique M. A. Verstegen and Luc J. W. van der Laan (Department of Surgery, Erasmus MC Transplant Center – University Medical Center, Rotterdam, The Netherlands) for providing the CCA organoids. We also thank Sven Plath and the Mechanical Workshop of the Biological Faculty for engineering the printer modifications and Ryan Sarkar for helpful discussion and proof-reading this work. Medical ethical approval for the use of patient liver tumor biopsies for research purposes has been granted by the Medical Ethical Committee (METC) of the Erasmus Medical Center in Rotterdam, The Netherlands (MEC-2013-143). Patient-provided written informed consent and all methods were performed in accordance with the relevant guidelines and regulations.

## Conflict of Interest

The authors declare no conflict of interest.

## Notes

https://github.com/LouiseBreide/BreidebandSupportingInformation/blob/main/Breideband_3Dbioprinter_Supplementary.pdf

### Competing Interest Statement

The authors have declared no competing interest.

## References

[1] B. P. Chan, K. W. Leong, Eur. Spine J. 2008, 17, 467.

[2] M. Tang, J. N. Rich, S. Chen, M. Tang, S. Chen, J. N. Rich, Adv. Mater. 2021, 33, 2004776.

[3] D. Sundaramurthi, S. Rauf, C. A. E. Hauser, Int. J. Bioprinting 2016, 2, 9.

[4] M. T. Vurat, C. Ergun, A. E. Elçin, Y. M. Elçin, Adv. Exp. Med. Biol. 2020, 1249, 67.

[5] A. Mazzocchi, S. Soker, A. Skardal, Appl. Phys. Rev. 2019, 6, 011302.

[6] I. Matai, G. Kaur, A. Seyedsalehi, A. McClinton, C. T. Laurencin, Biomaterials 2020, 226, 119536.

[7] K. Herrmann, K. Jayne, Animal Experimentation: Working Towards a Paradigm Change, Brill, 2019.

[8] P. Bédard, S. Gauvin, K. Ferland, C. Caneparo, È. Pellerin, S. Chabaud, S. Bolduc, Bioengineering 2020, 7, 1.

[9] G. A. Van Norman, JACC Basic to Transl. Sci. 2019, 4, 845.

[10] J. C. Fontoura, C. Viezzer, F. G. dos Santos, R. A. Ligabue, R. Weinlich, R. D. Puga, D. Antonow, P. Severino, C. Bonorino, Mater. Sci. Eng. C. 2020, 107, 110264.

[11] A. Sundarakrishnan, Y. Chen, L. D. Black, B. B. Aldridge, D. L. Kaplan, Adv. Drug Deliv. Rev. 2018, 129, 78.

[12] H. L. Ashe, J. Briscoe, Development 2006, 133, 385.

[13] H. B. Frieboes, X. Zheng, C. H. Sun, B. Tromberg, R. Gatenby, V. Cristini, Cancer Res. 2006, 66, 1597.

[14] J. Friedrich, C. Seidel, R. Ebner, L. A. Kunz-Schughart, Nat. Protoc. 2009, 4, 309.

[15] F. Bonnier, M. E. Keating, T. P. Wróbel, K. Majzner, M. Baranska, A. Garcia-Munoz, A. Blanco, H. J. Byrne, Toxicol. Vitr. 2015, 29, 124.

[16] D. Anton, H. Burckel, E. Josset, G. Noel, Int. J. Mol. Sci. 2015, 16, 5517.

[17] K. T. Lawlor, J. M. Vanslambrouck, J. W. Higgins, A. Chambon, K. Bishard, D. Arndt, P. X. Er, S. B. Wilson, S. E. Howden, K. S. Tan, F. Li, L. J. Hale, B. Shepherd, S. Pentoney, S. C. Presnell, A. E. Chen, M. H. Little, Nat. Mater. 2020, 20, 260.

[18] J. Gopinathan, I. Noh, Biomater. Res. 2018, 22, 11.

[19] A. S. Klar, S. Güven, T. Biedermann, J. Luginbühl, S. Böttcher-Haberzeth, C. Meuli-Simmen, M. Meuli, I. Martin, A. Scherberich, E. Reichmann, Biomaterials 2014, 35, 5065.

[20] A. Tong, Q. L. Pham, P. Abatemarco, A. Mathew, D. Gupta, S. Iyer, R. Voronov, SLAS Technol. 2021, 26, 333.

[21] J. Mielczarek, G. Gazdowicz, J. Kramarz, P. Łątka, M. Krzykawski, A. Miroszewski, P. Pieczarko, R. Szczelina, P. Warchoł, S. Wróbel, Solid State Phenom. 2015, 237, 221.

[22] Z. Wang, R. Abdulla, B. Parker, R. Samanipour, S. Ghosh, K. Kim, Biofabrication 2015, 7, 045009.

[23] T. A. Goldstein, C. J. Epstein, J. Schwartz, A. Krush, D. J. Lagalante, K. P. Mercadante, D. Zeltsman, L. P. Smith, D. A. Grande, Tissue Eng. 2016, 22, 1071.

[24] J. A. Reid, P. A. Mollica, G. D. Johnson, R. C. Ogle, R. D. Bruno, P. C. Sachs, Biofabrication 2016, 8, 025017.

[25] K. D. Roehm, S. V. Madihally, Biofabrication 2017, 10, 015002.

[26] D. T. Schmieden, S. J. Basalo Vázquez, H. Sangüesa, M. Van Der Does, T. Idema, A. S. Meyer, ACS Synth. Biol. 2018, 7, 1328.

[27] N. Bessler, D. Ogiermann, M. B. Buchholz, A. Santel, J. Heidenreich, R. Ahmmed, H. Zaehres, B. Brand-Saberi, HardwareX 2019, 6, e00069.

[28] M. Kahl, M. Gertig, P. Hoyer, O. Friedrich, D. F. Gilbert, Front. Bioeng. Biotechnol. 2019, 7, 184.

[29] B. Yenilmez, M. Temirel, S. Knowlton, E. Lepowsky, S. Tasoglu, Bioprinting 2019, 13, e00044.

[30] K. Ioannidis, R. I. Danalatos, S. Champeris Tsaniras, K. Kaplani, G. Lokka, A. Kanellou, D. J. Papachristou, G. Bokias, Z. Lygerou, S. Taraviras, Front. Bioeng. Biotechnol. 2020, 8, 1279.

[31] S. Agarwal, S. Saha, V. K. Balla, A. Pal, A. Barui, S. Bodhak, Front. Mech. Eng. 2020, 0, 90.

[32] S. Han, C. Min Kim, S. Jin, T. Young Kim, Biofabrication 2021, 13, 035048.

[33] Z. Wang, H. Kumar, Z. Tian, X. Jin, J. F. Holzman, F. Menard, K. Kim, ACS Appl. Mater. Interfaces 2018, 10, 26859.

[34] J. R. Yu, J. Navarro, J. C. Coburn, B. Mahadik, J. Molnar, J. H. Holmes, A. J. Nam, J. P. Fisher, Adv. Healthc. Mater. 2019, 8, 1801471.

[35] “DIY CO2 Incubator Bioreactor for Mammalian Cell Culture | Open Source Toolkit,” can be found under https://amchagas.github.io/open-source-toolkit/post/diy_co2_incubator_bioreactor_for_mammalian_cell_culture__pelling_lab/, accessed: December, 2021.

[36] S. Pedron, B. A. C. Harley, J. Biomed. Mater. Res. 2013, 101, 3404.

[37] X. Yu, T. Zhang, Y. Li, Polym. 2020, 12, 1637.

[38] Y. Wang, M. Ma, J. Wang, W. Zhang, W. Lu, Y. Gao, B. Zhang, Y. Guo, Materials (Basel). 2018, 11, 1345.

[39] P. Datta, M. Dey, Z. Ataie, D. Unutmaz, I. T. Ozbolat, npj Precis. Oncol. 2020, 4, 1.

[40] Y. L. Tsai, P. Theato, C. F. Huang, S. hui Hsu, Appl. Mater. Today 2020, 20, 100778.

[41] D. B. Kolesky, K. A. Homan, M. A. Skylar-Scott, J. A. Lewis, Proc. Natl. Acad. Sci. U.S.A. 2016, 113, 3179.

[42] J. M. Lee, W. L. Ng, W. Y. Yeong, Appl. Phys. Rev. 2019, 6, 011307.

[43] S. V Murphy, A. Atala, Nat. Biotechnol. 2014, 32, 773.

[44] J. A. Belgodere, C. T. King, J. B. Bursavich, M. E. Burow, E. C. Martin, J. P. Jung, Front. Bioeng. Biotechnol. 2018, 6, 66.

[45] B. S. Kim, G. Gao, J. Y. Kim, D. Cho, Adv. Healthc. Mater. 2019, 8, 1801019.

[46] M. Peter, A. Singh, K. Mohankumar, R. Jeenger, P. A. Joge, M. M. Gatne, P. Tayalia, ACS Appl. Bio Mater. 2019, 2, 916.

[47] L. S. S. M. Magalhães, F. E. P. Santos, C. de Maria Vaz Elias, S. Afewerki, G. F. Sousa, A. S. A. Furtado, F. R. Marciano, A. O. Lobo, J. Funct. Biomater. 2020, 11, 12.

[48] K. R. Mamaghani, S. M. Naghib, A. Zahedi, M. Rahmanian, M. Mozafari, Mater. Today Proc. 2018, 5, 15790.

[49] C. W. Macosko, Rheology: Principles, Measurements, and Applications, Wiley, 1996.

[50] M. Yin, J. Woollard, X. Wang, V. E. Torres, P. C. Harris, C. J. Ward, K. J. Glaser, A. Manduca, R. L. Ehman, Magn. Reson. Med. 2007, 58, 346.

[51] P. Kumar, T. Smith, R. Raeman, D. M. Chopyk, H. Brink, Y. Liu, T. Sulchek, F. A. Anania, J. Biol. Chem. 2018, 293, 12781.

[52] A. Charlesby, Int. J. Radiat. Appl. Instrumentation. Part C. Radiat. Phys. Chem. 1992, 40, 117.

[53] A. E. Rodríguez-Fraticelli, F. Martín-Belmonte, Methods Cell Biol. 2013, 118, 105.

[54] C. T. Gomillion, K. J. L. Burg, Compr. Biomater. II 2017, 6, 403.

[55] H. Mao, Y. Ito, Chapter 28 - Engineering Niches for Embryonic and Induced Pluripotent Stem Cells, Academic Press, 2017.

[56] S. P. Tarassoli, Z. M. Jessop, S. Kyle, I. S. Whitaker, Candidate Bioinks for 3D Bioprinting Soft Tissue, Woodhead Publishing, 2018.

[57] M. H. Zaman, L. M. Trapani, A. Siemeski, D. MacKellar, H. Gong, R. D. Kamm, A. Wells, D. A. Lauffenburger, P. Matsudaira, Proc. Natl. Acad. Sci. U. S. A. 2006, 103, 10889.

[58] E. J. Semler, C. S. Ranucci, P. V Moghe, Biotechnol. Bioeng. 2000, 69, 359.

[59] H. Holback, Y. Yeo, K. Park, in Biomed. Hydrogels (Ed.: S. Rimmer), Woodhead Publishing, 2011, pp. 3–24.

[60] W.-C. Yeh, P.-C. Li, Y.-M. Jeng, H.-C. Hsu, P.-L. Kuo, M.-L. Li, P.-M. Yang, P. H. Lee, Ultrasound Med. & Biol. 2002, 28, 467.

[61] A. Al-Abboodi, J. Fu, P. M. Doran, P. P. Y. Chan, Biotechnol. Bioeng. 2013, 110, 318.

[62] L. Broutier, G. Mastrogiovanni, M. M. A. Verstegen, H. E. Francies, L. M. Gavarró, C. R. Bradshaw, G. E. Allen, R. Arnes-Benito, O. Sidorova, M. P. Gaspersz, N. Georgakopoulos, B. K. Koo, S. Dietmann, S. E. Davies, R. K. Praseedom, R. Lieshout, J. N. M. IJzermans, S. J. Wigmore, K. Saeb-Parsy, M. J. Garnett, L. J. W. Van Der Laan, M. Huch, Nat. Med. 2017, 23, 1424.

[63] K. P. Lawrence, T. Douki, R. P. E. Sarkany, S. Acker, B. Herzog, A. R. Young, Sci. Rep. 2018, 8, 12722.

[64] D. Y. Wong, T. Ranganath, A. M. Kasko, PLoS One 2015, 10, 9.

[65] R. H. Utama, L. Atapattu, A. P. O’, K. Gaus, M. Kavallaris, J. J. Gooding, iScience 2020, 23, 101621.

[66] L. T. Li, G. Jiang, Q. Chen, J. N. Zheng, Mol. Med. Rep. 2015, 11, 1566.

[67] C. Widmann, in XPharm Compr. Pharmacol. Ref. (Eds.: S.J. Enna, D.B. Bylund), Elsevier, 2007, pp. 1–9.

[68] H. P. Sinn, A. Schneeweiss, M. Keller, K. Schlombs, M. Laible, J. Seitz, S. Lakis, E. Veltrup, P. Altevogt, S. Eidt, R. M. Wirtz, F. Marmé, BMC Cancer 2017, 17, 1.

[69] A. Kreso, J. E. Dick, Cell Stem Cell 2014, 14, 275.

[70] X. Yuan, J. Li, C. Coulouarn, T. Lin, L. Sulpice, D. Bergeat, C. De La Torre, R. Liebe, M. Gretz, M. P. A. Ebert, S. Dooley, H. L. Weng, Br. J. Cancer 2018, 119, 1358.

[71] K. Kawasaki, S. Kuboki, K. Furukawa, T. Takayashiki, S. Takano, M. Ohtsuka, Liver Int. 2021, 41, 865.

[72] P. Wang, L. Lv, P. Wang, L. Lv, Oncotarget 2016, 7, 81367.

[73] R. Kraiklang, C. Pairojkul, N. Khuntikeo, K. Imtawil, S. Wongkham, C. Wongkham, PLoS One 2014, 9, e89337.

[74] Pavel V. Korita, Toshifumi Wakai, Yoichi Ajioka, Makoto Inoue, Masaaki Takamura, Yoshio Shirai, Katsuyoshi Hatakeyama, Anticancer Res. 2010, 30, 2279.

[75] S. Dupont, L. Morsut, M. Aragona, E. Enzo, S. Giulitti, M. Cordenonsi, F. Zanconato, J. Le Digabel, M. Forcato, S. Bicciato, N. Elvassore, S. Piccolo, Nat. 2011, 474, 179.

[76] D. Loessner, C. Meinert, E. Kaemmerer, L. C. Martine, K. Yue, P. A. Levett, T. J. Klein, F. P. W. Melchels, A. Khademhosseini, D. W. Hutmacher, Nat. Protoc. 2016, 11, 727.

[77] “Particle Analysis,” https://imagej.net/imaging/particle-analysis, accessed: December, 2021.

[78] N. Köster, A. Schmiermund, S. Grubelnig, J. Leber, F. Ehlicke, P. Czermak, D. Salzig, Tissue Eng. 2016, 22, 552.

[79] J. Ye, G. Coulouris, I. Zaretskaya, I. Cutcutache, S. Rozen, T. L. Madden, BMC Bioinformatics 2012, 13, 134.

[80] J. D. Hunter, Comput. Sci. Eng. 2007, 9, 90.

[81] W. McKinney, presented at Proc. 9th Python Sci. Conf., June, 2010.

[82] M. Waskom, J. Open Source Softw. 2021, 6, 3021.

[83] N. A. Hotaling, K. Bharti, H. Kriel, C. G. Simon, Biomaterials 2015, 61, 327.

