## Supporting Information for "Bioprinting the Tumor Microenvironment with an Upgraded Consumer Stereolithographic 3D Printer"

**Figure S1**: CAD files for the modifications on the printer available on GitHub.

https://github.com/LouiseBreide/BreidebandSupportingInformation/blob/main/Bioprinting_8xIbidi_plate-holder.stl

https://github.com/LouiseBreide/BreidebandSupportingInformation/blob/main/Bioprinter_8xPlatform_Template.stl

https://github.com/LouiseBreide/BreidebandSupportingInformation/blob/main/Bioprinting_Support_platform_Ibidi.stl

https://github.com/LouiseBreide/BreidebandSupportingInformation/blob/main/Bioprinter_CompressedAirInlet.stl

**Figure S2**: Picture of the finished bioprinter after adaptations. The changes have been marked with red arrows.

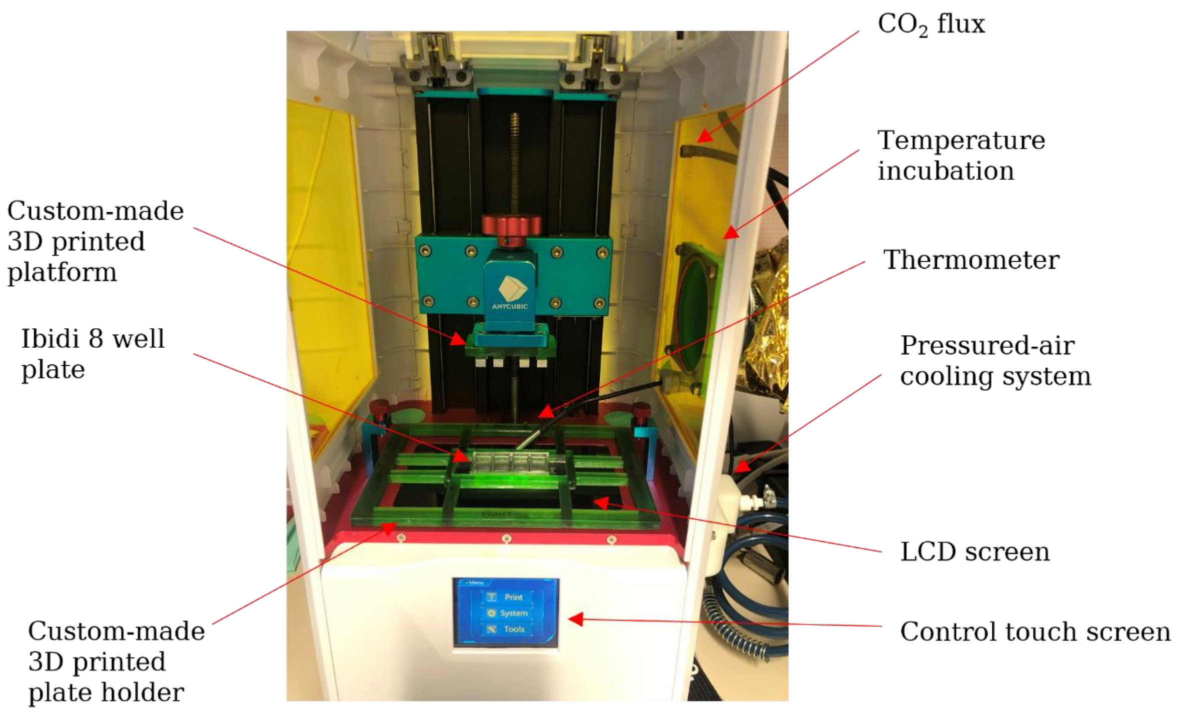


**Figure S3**: Complete measurements for the rheological analysis available on GitHub.

[https://github.com/LouiseBreide/BreidebandSupportingInformation/blob/main/Bioprinter_Summary_Rheology.xls](https://github.com/LouiseBreide/BreidebandSupportingInformation/blob/main/Bioprinter_Summary_Rheology.xls)

**Figure S4**: Cryo-FIB SEM picture of milling tests on a 3%/3% GelMA/PEGDA hydrogel sample. Squares (20 x 20 µm) were etched with increasing time to reveal deeper regions of the hydrogel and determine the optimal etching time.

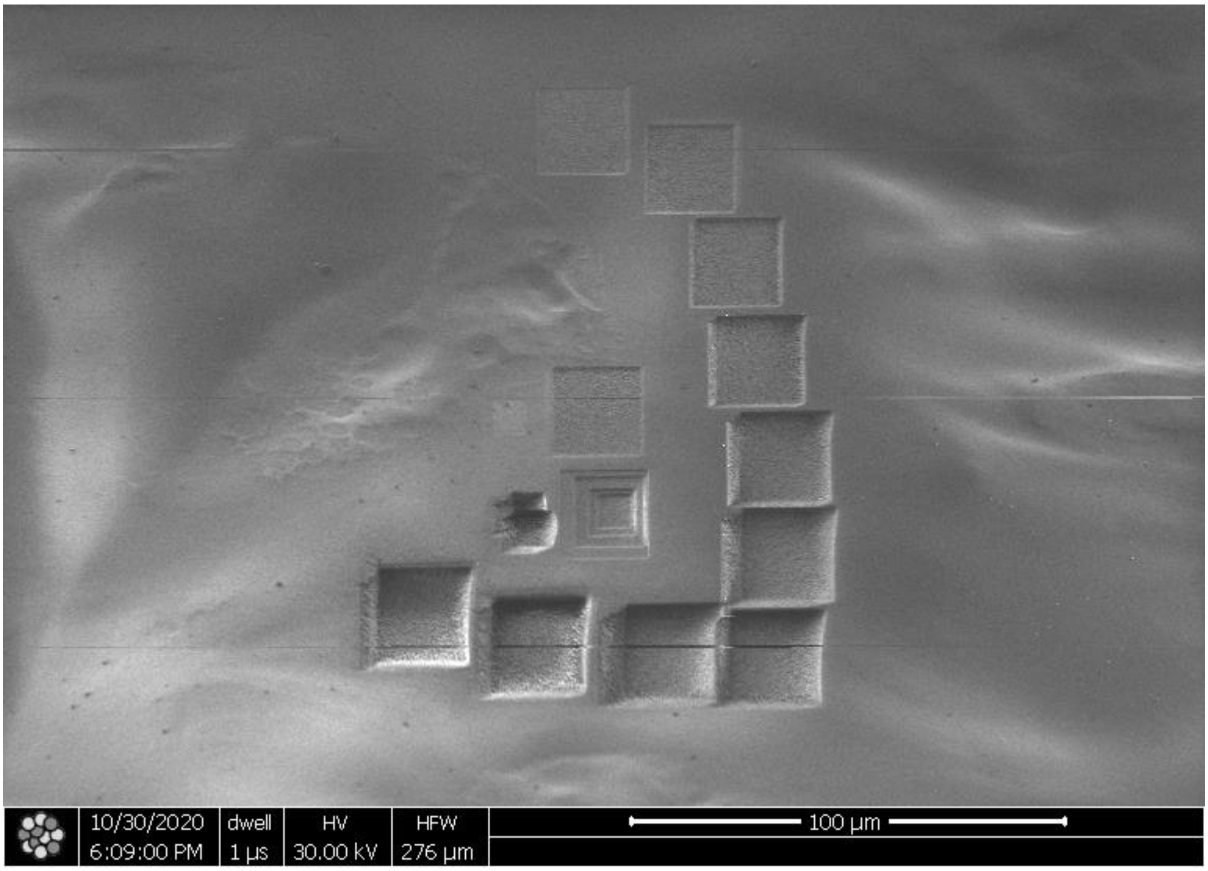


**Figure S5**: Imaging of CCA organoids displaying a compact morphology, taken at three different timepoints during a 7-day timelapse. As a writing convention, first the GelMA concentration was written, then the PEGDA concentration (for example, 3%/1.5% referred to 3% GelMA and 1.5% PEGDA). Microscope: Zeiss Axio Observer Z1. Objective: Plan-Apochromat 5x/0.16. Camera: AxioCam MR R3. Voxel size: 1.29 x 1.29 x 60 µm³. Scale bar: 100 µm.

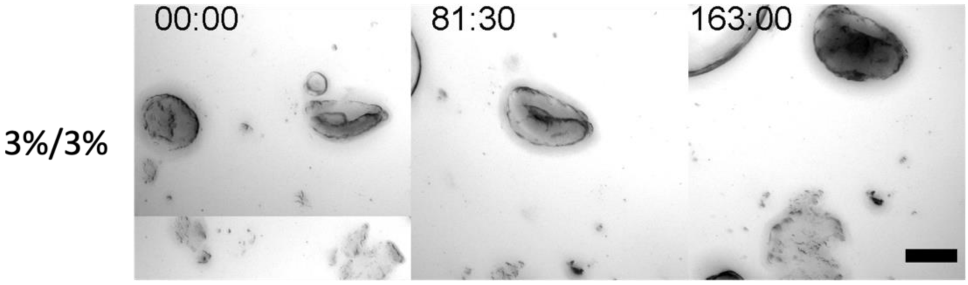


**Figure S6**: Cell viability of the CCA organoids in the bioprinted constructs after 7 days in culture.

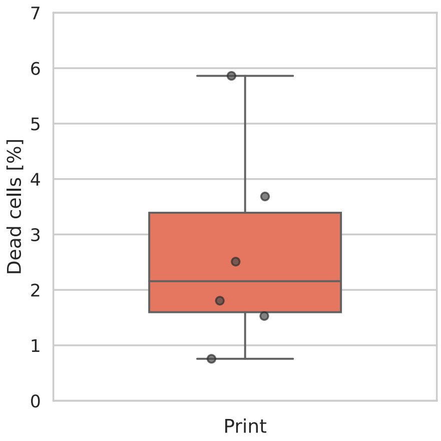


**Figure S7**: Complete RT-qPCR analysis available on GitHub.

https://github.com/LouiseBreide/BreidebandSupportingInformation/blob/main/Bioprinting_RT-qPCR_results.xlsx

**Figure S8**: Additional tumor-related markers were investigated. The variation in the results (upregulation versus downregulation) shows the heterogeneity of the tumor. The samples were normalized to the Matrigel samples. (A) Axin 2 (*AXIN2*) is part of the WNT signaling pathway which is a pharmacological target in cholangiocarcinoma treatments (Boulter et al. 2015, J Clin Invest.). *AXIN2* was found to be upregulated. *TNFRSF19* (TNF Receptor Superfamily Member 19) is part of the TGFβ pathway to promote tumorigenesis (Deng et al. 2017, Cancer Res.). *TNFRSF19* was downregulated. (B) Cystic fibrosis transmembrane conductance regulators (*CFTR)* are an indicator of high fibrosis in diseased tissues (Kim et al. 2002, Dig Dis Sci.). All samples were downregulated but the square one. Epithelial cell adhesion molecule (*EpCAM*) is a prognosis marker in cholangiocarcinoma (Sulpice et al. 2014, J Surg Res.). EpCAM was slightly downregulated compared to Matrigel samples.

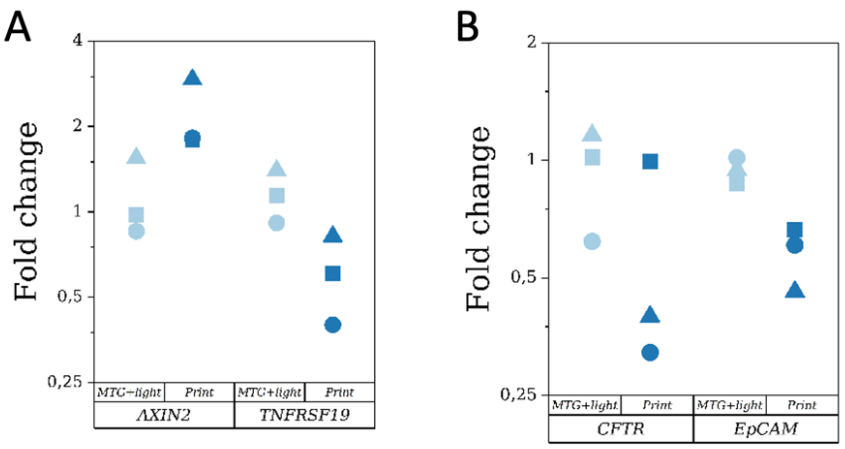


**Figure S9**: CAD of the platform for the Anycubic Photon S bioprinter for a 96 well-plate available on GitHub. https://github.com/LouiseBreide/BreidebandSupportingInformation/blob/main/96-well-plate_Anycubic-Bioprinter.stl

**Figure S10**: List of antibodies and dyes.

Table 1: List of primary antibodies

| **Antigen** | **Supplier** | **Cat. number** | **Clonality** | **Origin** | **Reactivity** | **Dilution** |
|---|---|---|---|---|---|---|
| Caspase 3 | Cell Signaling | 9661 | Polyclonal | Rabbit | Human, mouse, rat, monkey | 1:400 |
| Ki67 | Abcam | Ab6526 | Monoclonal | Rabbit | Human, mouse | 1:100 |
| Keratin 19 | St John's Laboratory | STJ24355 | Polyclonal | Rabbit | Human, mouse, rat | 1:100 |
| YAP | Santa Cruz Biotechnology | sc-101199 | Monoclonal | Mouse | Human, mouse, rat | 1:400 |

Table 2: List of secondary antibodies

| **Antigen** | **Supplier** | **Cat. number** | **Origin** | **Fluorophore** | **Dilution** |
|---|---|---|---|---|---|
| Rabbit | Thermo Fisher Scientific | A11008 | Goat | Alexa Fluor 488 | 1:400 |
| Rabbit | Thermo Fisher Scientific | A11011 | Goat | Alexa Fluor 546 | 1:400 |
| Mouse | Thermo Fisher Scientific | A21131 | Mouse | Alexa Fluor 488 | 1:400 |

Table 3: List of dyes

| Dye | Supplier | Cat. number | Dilution |
|---|---|---|---|
| Phalloidin 488 | Thermo Fisher Scientific | A12379 | 1:100 |
| Phalloidin 546 | Thermo Fisher Scientific | A22283 | 1:200 |
| Phalloidin 647 | Thermo Fisher Scientific | A22287 | 1:100 |
| Hoechst 33342 | Thermo Fisher Scientific | H1399 | 1:500 |

**Figure S11**: List of primers used for this work available on GitHub.

https://github.com/LouiseBreide/BreidebandSupportingInformation/blob/main/Bioprinter_RT-qPCR_primerslist.xlsx
